# Filtered finite state projection method for the analysis and estimation of stochastic biochemical reaction networks

**DOI:** 10.1101/2022.10.18.512737

**Authors:** Elena D’Ambrosio, Zhou Fang, Ankit Gupta, Sant Kumar, Mustafa Khammash

## Abstract

Time-lapse microscopy has become increasingly prevalent in biological experimentation, as it provides single-cell trajectories that unveil valuable insights into underlying networks and their stochastic dynamics. However, the limited availability of fluorescent reporters typically constrains tracking to only a few network species. Addressing this challenge, the dynamic estimation of hidden state-components becomes crucial, for which stochastic filtering presents a robust mathematical framework. Yet, the complexity of biological networks often renders direct solutions to the filtering equation intractable due to high dimensionality and nonlinear interactions.

In this study, we establish and rigorously prove the well-posedness of the filtering equation for the time-evolution of the conditional distribution of hidden species. Focusing on continuous-time, noise-free observations within a continuous-time discrete state-space Markov chain model, we develop the Filtered Finite State Projection (FFSP) method. This computational approach offers an approximated solution by truncating the hidden species’ state space, accompanied by computable error bounds. We illustrate the effectiveness of FFSP through diverse numerical examples, comparing it with established filtering techniques such as the Kalman filter, Extended Kalman filter, and particle filter. Finally, we show an application of our methodology with real time-lapse microscopy data. This work not only advances the application of stochastic filtering to biological systems but also contributes towards more accurate implementation of biomolecular feedback controllers.

**Author Summary:** The aim of this paper is to introduce a novel computational approach for numerically solving high-dimensional filtering problems associated with stochastic reaction network models in intracellular processes. This method, termed the Filtered Finite State Projection (FFSP) method, can reliably predict the dynamics of hidden species in reaction systems based on time-course measurements of the stochastic trajectories of certain species. While stochastic filtering is extensively utilised in engineering, its application in biology has been limited, primarily due to the nonlinear nature of biological interactions and the discrete, non-Gaussian nature of state variables. Traditional filtering techniques, such as the Kalman filter, often encounter difficulties under these conditions. We demonstrate that the FFSP method provides an accurate solution to the stochastic filtering problem, complete with a computable error bound. We present several numerical examples to showcase the effectiveness of FFSP and its superior performance compared to other filtering methodologies. Additionally, we apply FFSP to biological data, successfully reconstructing the hidden dynamics of a yeast transcription system from partial measurements obtained through time-lapse microscopy. We believe that FFSP could be a valuable tool for elucidating hidden intracellular dynamics and understanding stochastic cellular behaviours.

## 1 Introduction

The genetics revolution of the 20^th^ century significantly impacted various foundational aspects of biological sciences. A landmark event during this era was the discovery of Green Fluorescent Protein in 1962, which spurred the development of fluorescent technologies [1] and microscopic techniques [2]. These innovations greatly enhanced scientists’ ability to observe the dynamic behaviors of living cells. However, despite these technological advances, tracking capabilities are still limited to a few cellular components, such as fluorescent proteins and mRNAs [3–5]. Consequently, critical dynamic states like gene activation, transcription factors, and promoter activities remain elusive and poorly understood.

Moreover, the inherently stochastic interactions among biomolecular species within cells, especially at low concentrations, add layers of complexity to understanding these biological systems [6–8]. This stochasticity at the gene expression level introduces significant phenotypic variability even among genetically identical cells [9, 10], which is crucial for understanding cellular decision-making processes [11]. To fully characterize the stochastic behaviour in living cells, the last decades have seen a surge of development in stochastic models and forward simulation techniques in biology [12–14].

One of the most common ways to model stochastic biochemical reaction processes is to employ a continuous-time Markov Chain (CTMC) with discrete state space which keeps track of the molecular species copy number over time. The states’ probability of this Markov Chain satisfies a very large or possibly infinite dimensional linear ordinary differential system of coupled equations, also known as the Chemical Master Equation (CME), that describe the chain’s state probabilities over time. One well-known numerical strategy for solving this extensive system is the finite state projection (FSP) method [15], which truncates the Markov Chain’s state space and gives rise to a finite-dimensional system for the evolution of state probabilities while providing an exact error certificate of the approximated solution.

In addition to this relatively direct approach, the CME can also be solved by simulation based approaches, like the Gillespie algorithm [14, 16] and its variants [17, 18], which exactly simulate the Markov chain model and approximate the probability distribution with the empirical distribution. However, these simulation-based approaches become computationally infeasible when the system size increases or several reactions are firing in a small time unit. Therefore, other approximations of the CME have become available. Some of these methods, such as the Chemical Langevin Equation and the Linear Noise Approximation (see [19] for a detailed reference), focus on approximating the underlying stochastic process of the CME with a continuous state process. Instead, other methods look at the CME solution statistics and provide moment closure techniques to close the moment equations [20–22].

Despite the availability of several approaches to approximate the CME’s solution, comparatively fewer computational methods exist for obtaining real-time estimations of latent cellular states as the observations arrive sequentially in time [23–28]. Experimentally, we can only observe certain chemical species, making it challenging to understand the hidden parts of the network that generated the observed data. Understanding the behaviour of the hidden species in relation to observed data through network simulation is typically computationally infeasible, and an inordinately large number of forward stochastic simulations would need to be generated to obtain a sample of trajectories that is consistent with observations.

Thus, it is more practical to use observed data as it arrives to predict hidden dynamics. Stochastic filtering offers a viable and accurate approach for reconstructing and understanding stochastic reaction networks beyond current experimental reach. In particular, stochastic filtering theory offers a robust framework for real-time estimation of latent states from time-course measurement data comprising of partial state observations. Specifically, it involves estimating the conditional probability distribution *π*_*t*_ of the latent state, modeled by a stochastic process *X*_*t*_, based on all available information up to time *t*. This information is typically encapsulated in a sigma-algebra generated by an observation process, denoted as 𝒴_*t*_. The conditional probability *π*_*t*_ usually satisfies a complex stochastic evolution equation known as the filtering equation, which is generally non-linear and infinite-dimensional [29].

The complexity and high-dimensionality of this equation pose significant challenges for real-time estimation of latent cellular states from time-course measurement data. Consequently, developing computational methods that address these challenges and reduce the computational time required for accurately solving the filtering equation is essential. This would facilitate real-time estimations of hidden cellular states, allowing for more effective experimental design. With real-time estimations, we could further explore poorly understood cellular functions, analyze the impact of noise on cellular regulation, and improve the design of biomolecular controllers, which hold tremendous potential for therapeutic applications [30].

One of the first pivotal advances in stochastic filtering has been the linear filter introduced by Kalman in the discrete-time setting [31]. The extension to the continuous-time case was given later by Bucy [32]. For linear dynamical systems corrupted with Gaussian noise, this linear filter provides an analytical expression for the conditional distribution, *π*_*t*_, which is Gaussian and therefore determined by its mean and covariance matrix. This Kalman-Bucy filter has been widely used in practical applications, including space missions like the Apollo project [33]. For the non-linear setting, the filtering equation (referred to as the Kushner-Stratonovich equation [34, 35] in some cases) is generally infinite-dimensional [36]; an analytical expression of the solution is not always available. Facing this challenge, a surge of interest has occurred in particle filter methods since the seminal paper by Gordon, Salmond, and Smith [37]. The key feature of such methods is that finitely many random samples (or particles) are generated to represent the posterior probability distribution. These methods look promising and particularly suited to address the infinite-dimensional feature of filtering problems.

In the stochastic reaction networks framework, state and parameter estimations have been mainly carried out with the Kalman filter and particle filtering (also known as the sequential Monte-Carlo method). In [38], and [39], Sun et al. and Chuang et al. applied the Extended Kalman filter and the Kalman filter to non-linear state-space models of gene regulatory networks. Moreover, in [40], Calderazzo et al. used an Extended Kalman-Bucy filter to obtain parameters and state estimations in discrete-time of a stochastic negative feedback model. In [41], Liu et al. employ (i) extended Kalman filter (EKF), (ii) unscented Kalman filter (UKF) and (iii) the particle filter to estimate unobserved states of cellular responses in *Escherichia Coli* subjected to a sudden temperature increase. Moreover, in [42], and [43], Boys et al. and Golightly et al. applied sequential Monte-Carlo methods to diffusion approximations models of gene regulatory networks to infer kinetic parameters. Along these lines, Fang et al. [27, 28, 44] used a particle filter algorithm on a reduced model of a stochastic reaction network, derived through time-scale separation, rigorously showing that the filter of the original model converges to the one of the reduced model. In the context of noisy and discrete time observation processes, Huang et al. [25] derive the posterior master equation and approximate the infinite-dimensional moment dynamics, validating their method using both in silico data and single-cell data from two gene expression systems in *Saccharomyces cerevisiae*. For a noise-free observation process, in [45–47], Zechner et al. solved the filtering equation through a moment-closure approach. In the same setting, Rathinam et al. [26] proposed a bootstrap particle filtering (BPF) algorithm to solve numerous estimations problems in biologically relevant examples. In [23], Rathinam et al. develop a novel particle filtering method, called the targeting algorithm, to solve the stochastic filtering problem in the context of noise free discrete time observations.

The widespread adoption of traditional filtering techniques faces significant challenges due to their inherent limitations in scenarios in Biology that diverge from their core assumptions. The Kalman filter, which relies on assumptions of system linearity and Gaussian noise, often falls short in the complexities of real-world applications characterized by nonlinear dynamics and non-Gaussian disturbances. Consequently, the Extended Kalman Filter (EKF) was developed to improve modeling accuracy for nonlinear systems. However, both the Kalman filter and EKF exhibit difficulties in discrete-state processes with pronounced deviations from Gaussian noise, particularly in situations involving low copy numbers.

To address these challenges, we compared the performance of these conventional filters using a nonlinear chemical reaction network model with feedback (details in Section **S4** of the Supplementary Material), as depicted in panel (a) of Fig. 1. This comparison assesses their effectiveness relative to a bootstrap particle filter (BPF) developed in [26] and the Filtered Finite State Projection (FFSP) method, the latter being a contribution of this article. In panel (d) of Fig. 1, the SSA trajectory of the hidden processes (*Z*_1_ and *Z*_2_) is presented alongside the conditional mean and standard deviations estimated by the various filters.

**Fig. 1.**
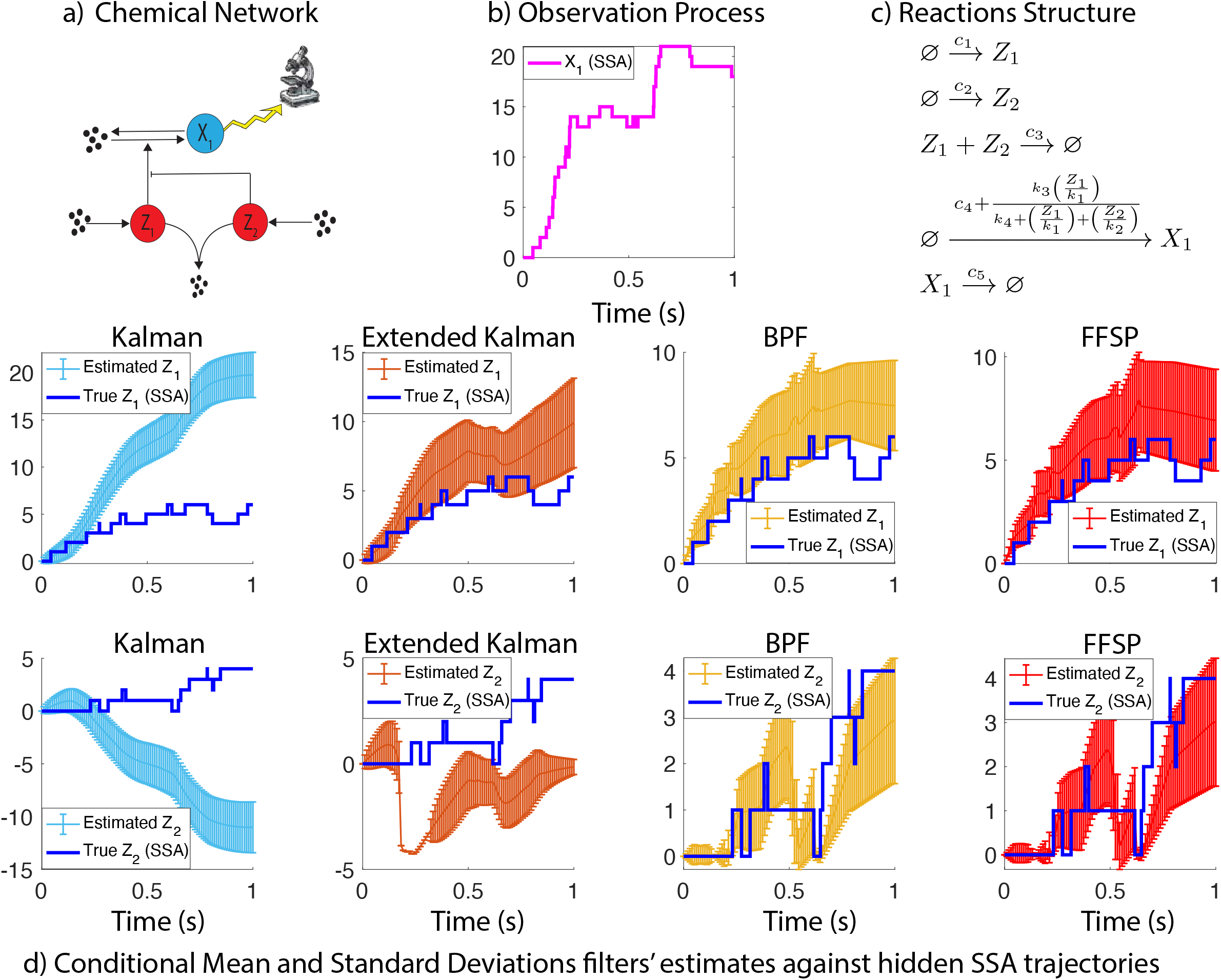
Non-Linear Network with Feedback. **a). Chemical Reaction Network Structure:** The hidden species *Z*_1_ and *Z*_2_, highlighted in red, are produced and jointly degraded. They also catalytically influence the production of *X*_1_, the observed species in the network. **b). Observation SSA Process Trajectory:** The trajectory of species *X*_1_ is fed as input to the filters in panel (d). **c). Chemical Reactions Structure:** The network’s chemical reactions are structured with the following parameters: *c*_1_ = 10, *c*_2_ = 10, *c*_3_ = 0.5, *c*_4_ = 0, *c*_5_ = 0.5, *k*_1_ = 1000, *k*_2_ = 1, *k*_3_ = 1000, *k*_4_ = 0.04. The initial condition is [*z*_10_, *z*_20_, *x*_10_]^*T*^ = [0, 0, 0]^*T*^. **d). Filter Performance and Estimations:** The Kalman and Extended Kalman filters, Bootstrap Particle Filter (BPF), and FFSP conditional expectations and variance estimations are plotted against the exact hidden trajectories of *Z*_1_ and *Z*_2_, corresponding to the observation process trajectory shown in panel (b).

Given the observation process trajectory of *X*_1_ shown in panel (b) and the feedback structure presented in panel (c), it appears that the Kalman filter may struggle to accurately reconstruct the behavior of the hidden species. This is suggested by the fact that the trajectories of the hidden processes fall outside the one-interval standard deviation bands. In particular, the Kalman filter seems to have difficulty accounting for the roles of *Z*_1_ and *Z*_2_ in the initial sharp increase of *X*_1_, possibly due to the highly nonlinear nature of the feedback.

However, the Extended Kalman filter, designed to overcome the nonlinearities inherent in the network, shows better performance in estimating the hidden species’ behavior, though a significant mismatch is observed for the hidden species *Z*_2_. Specifically, the Extended Kalman filter provides negative estimates, likely due to a mismatch between the observation models. The EKF expects Gaussian-like observation noise suited to continuous state processes that can take both positive and negative values, while the Poisson-like noise in this context is designed for non-negative discrete state processes. Therefore, in the presence of strong nonlinearities and non-Gaussian noise, both the Kalman and Extended Kalman filters may not be suitable for predicting the hidden dynamics of discrete state processes.

In stark contrast, the particle filter and the FFSP method exhibit accuracy in predicting the hidden species’ behavior, affirming their effectiveness for nonlinear discrete stochastic systems affected by non-Gaussian noise. The FFSP, in particular, offers a valid alternative for the analysis of biological systems modeled as continuous-time Markov chains (CTMC).

However, as it can be seen in Section 3.1.1, Kalman filter can still be a valuable resource for state estimation in scenarios of linear networks, no feedback, and an observation process with noise that approximates Gaussian noise close enough.

Moreover, while particle filters are known for their precision in estimating summary statistics of conditional distributions, their efficacy significantly diminishes when calculating specific event probabilities, especially for rare events. Our example, as highlighted in Fig. 2, demonstrates that the bootstrap particle filter’s relative error [26] in estimating the low-probability event of the system being in state 0 in a very simple linear network remains substantial, irrespective of an increase in particle numbers, which consequently escalates computational demands. This observation underscores a critical limitation of particle filters, emphasizing the need for more sophisticated methodologies in scenarios dominated by rare events.

**Fig. 2.**
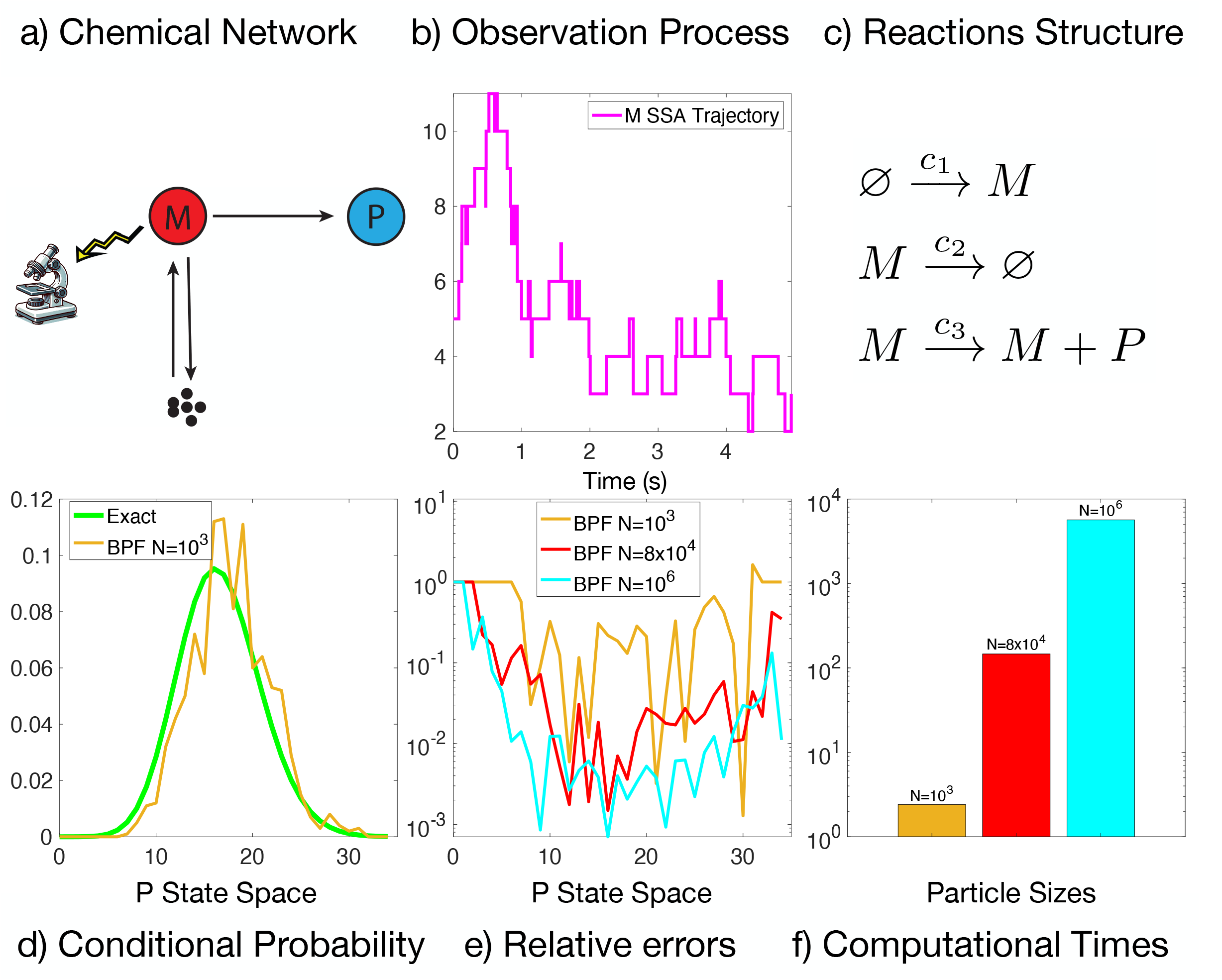
Transcription-Translation Network. **a). Chemical Reaction Network Structure**: The hidden species *P*, highlighted in blue, is being translated by the observed species *M*, highlighted in red, which in turn is being produced and degraded. **b). Observation SSA Process Trajectory**: The trajectory of species *M* for which the conditional distribution and the relative errors have been computed in panel (d). **c). Chemical Reactions Structure**: The network’s chemical reactions are structured with the following parameters: *c*_1_ = 1, *c*_2_ = 5, *c*_3_ = 1 and initial conditions [*p*_0_, *m*_0_]^*T*^ = [0, 5]^*T*^ **d). Conditional Distribution Comparison**: Estimation of the conditional distribution of the network for the trajectory given in panel (b) computed with a bootstrap particle filtering (BPF) with *N* = 1000 particles and compared with the exact analytic expression. **e). Relative Errors** for different particle sizes of the *L*_1_ difference of the exact and estimated conditional distributions. **f). Computational Times** for the different particle filters sizes.

This discussion emphasizes the distinct challenges encountered by the Kalman, Extended Kalman, and particle filters and it also highlights why the FFSP method could be a valid alternative for modeling and estimation in complex biological systems.

In this paper, we concern ourselves with the situation of noise-free observations, which is reasonable given the significant advances in modern microscopy. In this setting, we first rigorously verified the validity of the continuous-time filtering equation, previously derived in [26, 45, 46]. The filtering equation has an unnormalized version, which resembles the CME very closely except for a jump term driven by the observation process. Inspired by this, we propose the Filtered Finite State Projection (FFSP) approach for the filtering problem by solving the unnormalized filtering equation on the truncated state space of the hidden species. Since the FFSP method solves the filtering equation in a direct fashion, it turns out to be more accurate than simulation-based particle filtering. We provide two different equivalent versions of the FFSP algorithm together with their error bounds. Finally, several biologically relevant numerical examples are presented to illustrate our approach.

The paper is organised as follows. First, we introduce our methods in the next section. Specifically, in Section 2.1 and Section 2.2, we briefly review basic concepts of stochastic reaction networks and the associated stochastic filtering problem in the situation of continuous-time noise-free observations. In Section 2.3 we outline the Filtered Finite State Projection (FFSP) method together with its accuracy analysis, and in Section 2.4, we propose an alternative FFSP algorithm with a tighter error bound but with more limited applicability. Then, in Section 3 we present several numerical examples in which we compare the performance of both the algorithms, compute the error bounds, and compare our algorithm with the bootstrap particle filter (BPF) algorithm available in [26] and the famous Kalman filter. In Section 3.4, we evaluated our method using the telegraph model within a hybrid experimental framework, as detailed in [48]. Specifically, the transcription component of the circuit was operationalized in yeast cells [49], with mRNA molecules marked by fluorescent reporters. These trajectories were subsequently utilized to synthesize in silico SSA protein translation profiles. We employed these protein dynamics as observational data to infer mRNA trajectories using the FFSP. Our approach demonstrated high accuracy and closely aligned with the biological data, underscoring its effectiveness in practical biological applications. Finally, Section 4 concludes this paper.

### Notations

In this paper, we term **0** as a zero vector of a proper size and 𝟙(·) as the indicator function, which equals one if the argument holds and zero otherwise. **1** is a vector of ones of a proper size.

## 2 Materials and methods

### 2.1 Stochastic modelling of intracellular chemical reacting processes and their associated filtering problems

We consider an intracellular chemical reacting system that has *n* species (*S*_1_, …, *S*_*n*_) and *M* reactions:

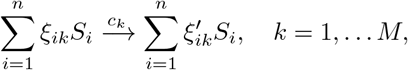

where *ξ*_*ik*_ and 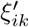 are the numbers of *S*_*i*_ molecules consumed and produced in the *k*^th^ reaction. We indicate with *a*_1_, *a*_2_, … *a*_*M*_ the propensity functions representing the rates of these *M* reactions. Moreover, we term 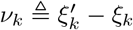, as the stoichiometry vector associated with the *k*^*th*^ reaction channel and define the stoichiometry matrix as follows:

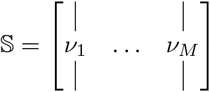

In the low copy number regime, the nature of the interactions among intracellular biomolecular species is inevitably stochastic, and its dynamics is usually modelled by a stochastic dynamic equation, known as the Random Time Change (RTC) representation [12]:

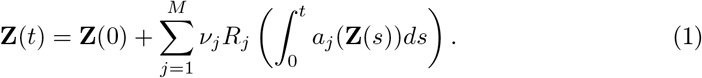

Here, **Z**(*t*) is a continuous-time discrete state Markov Chain keeping track of the molecules copy numbers, and *R*_1_, …, *R*_*M*_ are independent unit rate Poisson processes, which count the firing events of every reaction. We only consider the processes **Z**(*t*) that satisfy the following non-explosivity condition:

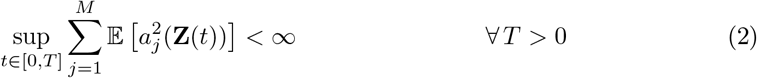

which suggests that the system state will almost surely not grow to infinity in a finite time. Also, to avoid negative molecular copies, we assume that *a*_*j*_(*z*) = 0 for every *j* ∈ {1, …, *M*} and 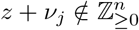.

The probability distribution to the process in (1) is given by the Kolmogorov’s forward equation also known as the Chemical Master Equation (CME) [12]:

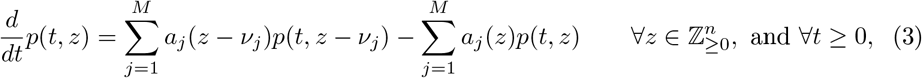

where *p*(*t, z*) = *P* {**Z**(*t*) = *z*} is the (unconditional) probability mass function associated with the Markov Chain, indicating the probability of finding the chemical system in state *z* ∈ ℤ^*n*^ at time *t >* 0. Under the condition (2), the two dynamical representations (1) and (3) of the system are equivalent.

With modern time-lapse fluorescent microscopes, scientists can now measure single-cell trajectories at high resolutions [1, 2]. However, due to the limited choices of distinguishable fluorescent reporters, this technology cannot keep track of all the chemical species in a cell [3]. This obstacle gives rise to a stochastic filtering problem for the considered system, i.e., to infer the hidden state based on the partially observed cell states. An accurate solution to this problem can provide deep insights into intracellular dynamics and enables the design of better controllers [30].

### 2.2 The Filtering Problem for the Noise-Free Observation Process

To describe this filtering problem, we further decompose the system into two sub-networks, **Z**(*t*) = (**X**(*t*), **Y**(*t*)), where 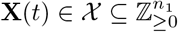 keeps track of the copy numbers of the *n*_1_ hidden (unobserved) species, and 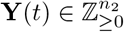(with *n*_2_ = *n* − *n*_1_) keeps track of the copy numbers of the observed species. For simplicity, we rearrange the species orders so that the first *n*_1_ species are the hidden species, and the last *n*_2_ species are the observed ones. In this paper, we assume that we can time-continuously and exactly (noise-freely) observe the trajectory of the observed species **Y**(*t*). Our goal is to compute the conditional distribution *π*(*t, x*) ≜ *P*{**X**(*t*) = *x*|**Y**(*s*) = *y*(*s*), 0 ≤ *s* ≤ *t*}^1^, which is also viewed as the optimal estimate of the hidden process given the observations because for any 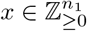 and *t* ≥ 0

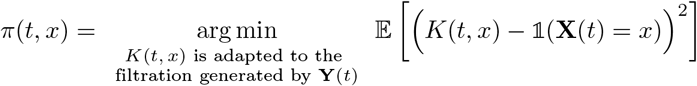

where adapted means that the process under consideration is measurable with respect to the filtration generated by **Y**(*t*) for every *t*≥ 0.

To present the filtering equation characterising the time evolution of *π*(*t, x*), we need first to introduce several notations related to the system decomposition. For each reaction stoichiometric vector *ν*_*j*_ (*j* = 1, …, *M*), we denote its first *n*_1_ components by 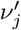 and the last *n*_2_ components as 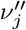. Following the notations in [26], we term 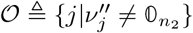 as the set of observable reactions that can alter the dynamics of **Y**(*t*), and we term, 𝒰 ≜ 𝒪^*c*^ as the unobservable reactions that cannot alter **Y**(*t*). We define *a* ^𝒪^(*x,y*) ≜Σ_*j* ∈𝒪_ *a*_*j*_(*x, y*), which is the total propensity of the observed reactions. Also, we term 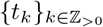 as the random jump times of the observed process **Y**(*t*) and denote 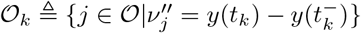(for every *k* ∈ ℤ_*>*0_), which represents the reaction channels compatible with the jump of the observation process at time *t*_*k*_. Finally, we define *t*_0_ = 0.

The filtering equation for the conditional distribution *π*(*t, x*) has been formally derived in [50] for systems with finite state-spaces. In the infinite state-space setting, instead, it has been informally obtained by [26, 45, 46], and it is expressed by

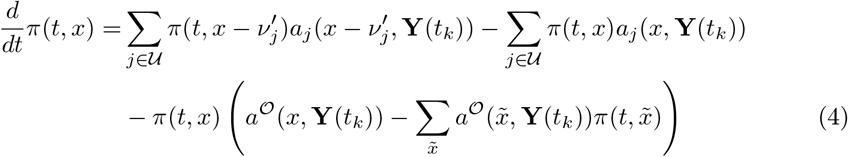

for all *k* ∈ ℤ_≥0_, *t* ∈ (*t*_*k*_, *t*_*k*+1_), and 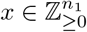, and

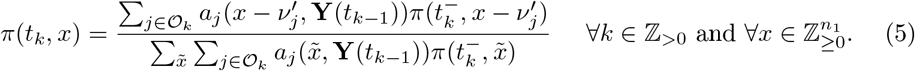

#### Theorem 1.

*Under the non-explosivity assumption* (2), *the conditional distribution π*(*t, x*) *is the unique (up to indistinguishability) non-negative solution of the filtering equation* (4) *and* (5) *that starts from π*(0, ·) *and satisfies*

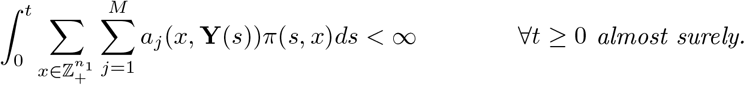

The proof of this theorem is presented in Section **S.1.3** of the supplementary material, where it is structured into two distinct parts: initially proving the validity of the filtering equation, followed by demonstrating the uniqueness of the solution. The filtering equation (4) is nonlinear due to the last term (being quadratic in *π*(*t, x*)), which brings many difficulties to solving this equation. To tackle this issue, [26] provided a Monte-Carlo method based on a linear equation characterising the un-normalised conditional probability *ρ*(*t, x*), from which you can obtain the filtering distribution via normalisation. Specifically, this un-normalised distribution is given by 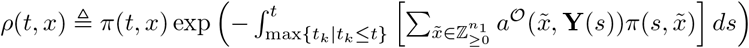 and therefore satisfies

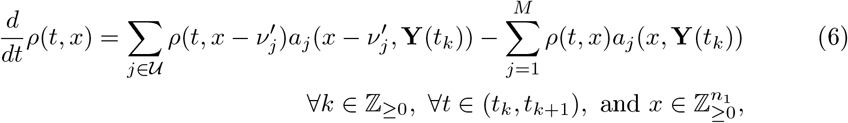

and

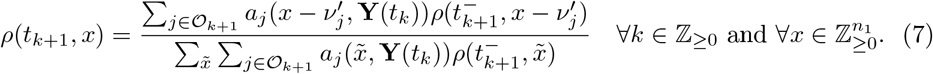

Then, the authors in [26] related the linear equation (6) with a new chemical reacting system and a random variable associated with that; afterward, a particle filter (a type of Monte-Carlo method) was proposed to compute the filtering equation. Different from this Monte-Carlo method, the literature [25, 45, 46] proposed moment closure for the filtering equation (4) and (5) to compute the conditional moments of *π*(*t, x*). More recently, an autonomous algorithm for this moment closure was proposed in [47].

### 2.3 Filtered finite state projection method

Here, we introduce a new method for the aforementioned filtering problem using the finite state projection (FSP) approach proposed in [15]. The idea is that the filtering equation (4) has a close resemblance to the CME (3), and, therefore, it is reasonable to expect that the FSP (an effective approach for solving CME) would be applicable to this filtering problem after some extensions. In this paper, we name this new method for the filtering problem as filtered finite state projection (FFSP) method.

Similar to the FSP which solves the CME directly by truncating the system’s state space, the FFSP also straightforwardly solves the linear differential equation (6) in the time intervals (*t*_*k*_, *t*_*k*+1_) (for *k* ∈ ℤ_>0_) by truncating the hidden species’ state space and updates the filter according to (7) at the jump times 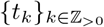. After approximating the un-normalised distribution *ρ*(*t, x*), the FFSP obtain estimates of *π*(*t, x*) by normalisation. To be more specific, when solving (6), the FFSP decomposes the hidden-state space 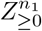 into a finite but large state space 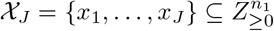 and an infinite-dimensional space 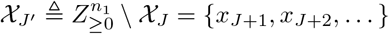 then the FFSP only solves (4) on the finite space 𝒳_*J*_. More precisely, by denoting *ρ*(*t*, 𝒳_*J*_) = (*ρ*(*t, x*_1_), …, *ρ*(*t, x*_*J*_))^⊤^ and *ρ*(*t*, 𝒳_*J*′_) = (*ρ*(*t, x*_*J*+1_), *ρ*(*t, x*_*J*+2_), …,)^⊤^, the linear equation (4) can be equivalently written as

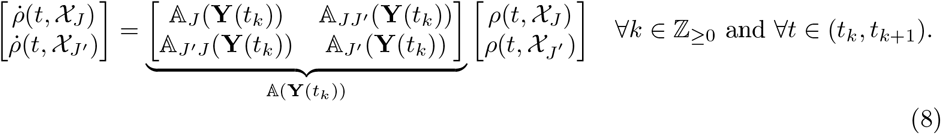

where 𝔸(**Y**(*t*_*k*_)) is an infinite dimensional matrix with each element defined by

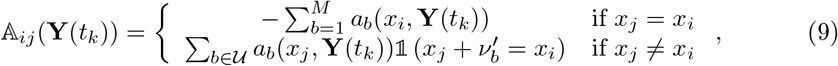

and 𝔸_*J*_ (**Y**(*t*_*k*_)), 𝔸_*JJ*′_ (**Y**(*t*_*k*_)), 𝔸_*JJ*′_ (**Y**(*t*_*k*_)), and 𝔸_*J*′_ (**Y**(*t*_*k*_)) are sub-matrices of A(**Y**(*t*_*k*_)) with proper dimensions. Instead of solving the infinite dimensional ODE (8), the FFSP only implements a finite dimensional ODE

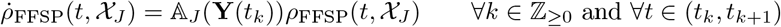

to approximate (8). Here, *ρ*_FFSP_(*t*, 𝒳_*J*_) is the FFSP’s estimates to the the un-normalised filter *ρ*(*t*, 𝒳_*J*_), and the solution of this finite dimensional ODE is

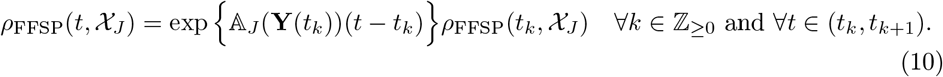

The detailed algorithm for the FFSP method is presented as Algorithm 1.

#### Algorithm 1 Filtered finite state projection (FFSP)

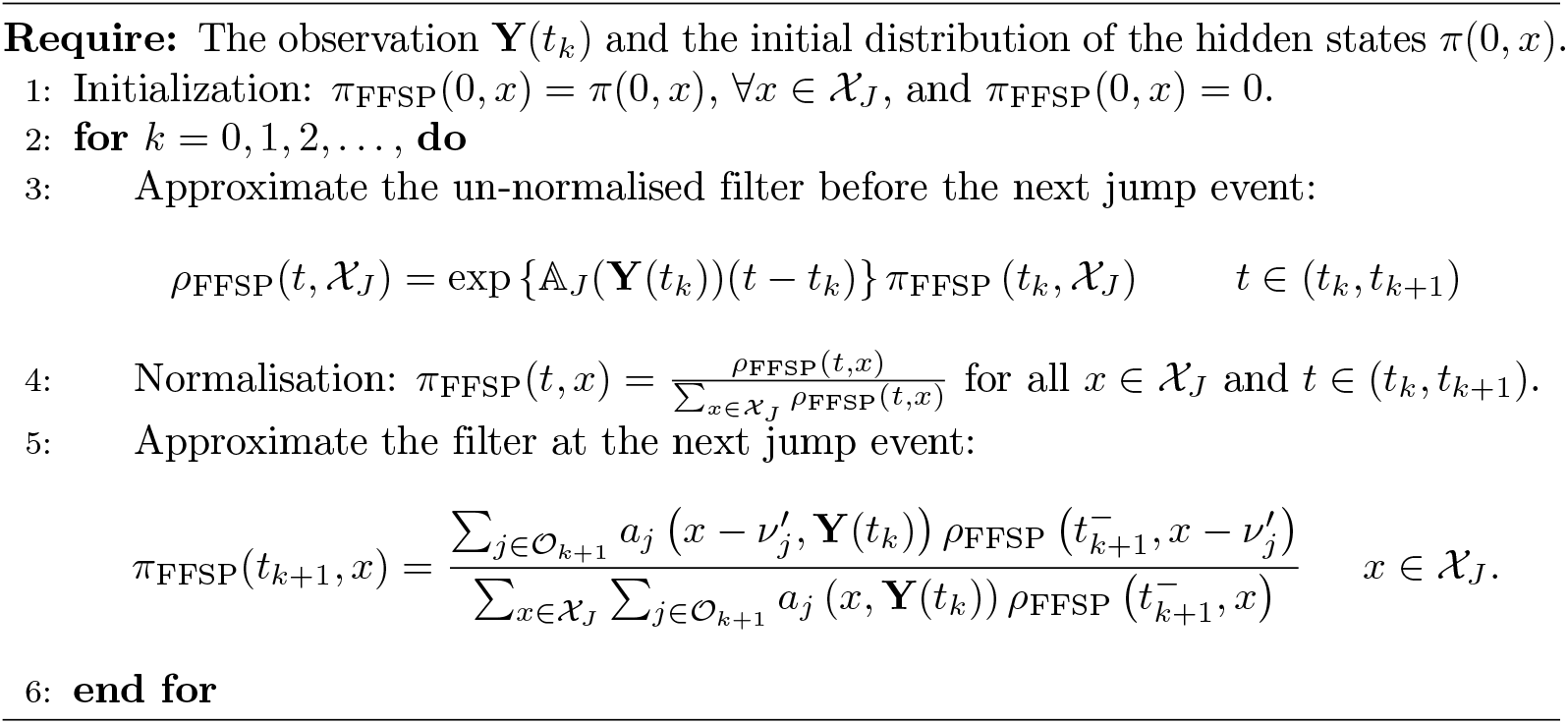

In general, both the original FSP approach and our FFSP method solve their associated problems in a straightforward manner, and they therefore can reach good accuracy at a potential cost of high computational complexity in large dimensional systems. However, they still have some major differences in their structures. For instance, compared with the original FSP [15] which only solves one single finite-dimensional ODE, our FFSP algorithm needs to compute the ODEs (10) recursively for each time interval (*t*_*k*_, *t*_*k*+1_) due to the recursive structure of the filtering equation. Furthermore, as can be seen in Algorithm 1, the approximated FFSP solution *π*_FFSP_(*t, x*) is not consistently component-wise upper bounded by the exact filter *π*(*t, x*) due to the normalization step. This phenomenon could be described as a ‘dominance property’ held by the exact solution, which is maintained in the original FSP algorithm and results in an exact error certification. However, in this context, the normalization procedures cause the loss of this property, making precise error bounds more challenging to establish.

Also, our approach has several advantages over other existing computational methods for solving the filtering equation (4) and (5). Compared with the moment-closure approach in [45, 46], our FFSP method provides more detailed information about the conditional distribution, e.g., the specific conditional probability of being in any subset of the state-space, in addition to the conditional moments. Moreover, as a Monte-Carlo method, the particle filtering approach can provide good estimates for summary statistics (e.g., conditional mean and variance) but less accurate estimates for specific conditional probabilities (especially for the rare events). In contrast, our method (as a direct approach) can consistently provide very accurate estimates of the conditional probability at each state. More detailed comparisons between the particle filtering and the FFSP are presented in Section 3.

Once the FFSP method is established, we are interested in the estimation error of this algorithm, which characterises the reliability of this method. In the following, we perform error analysis for the FFSP. For technical reasons, we only consider the case where the propensity function *a*^𝒪^(·, *y*) is upper bounded for each fixed 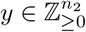, i.e.,

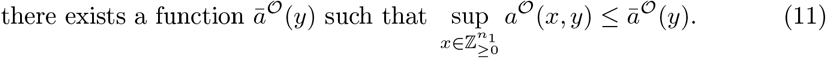

Moreover, we also assume that the network with only unobserved reactions

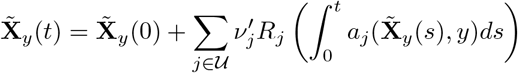

satisfies the conditions

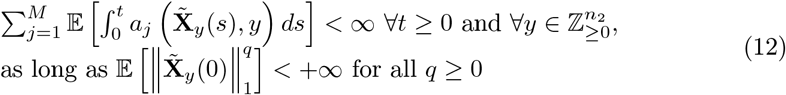

which guarantees the process to be non-explosive when initial conditions have finite moments. Both of these assumptions are reasonable and easily checkable in practice. Usually, under a microscope, scientists observe fluorescent reporters, which are catalytically produced and spontaneously degraded. In this setting, all the observable reactions satisfy (11) because of the saturation of catalytic production rates (due to the limited resources) and the independence of the degradation rate with other species. Also, some easily checkable conditions for the assumption (12) have already been provided in [51], and those conditions are shown to cover a large class of practical biochemical reacting systems.

An upper bound for the estimation error of Algorithm 1 is given in the following theorem.

#### Theorem 2.

*Let us denote filters π*_*FFSP*_(*t*, 𝒳) ≜ (*π*_*FFSP*_(*t, x*_1_), *π*_*FFSP*_(*t, x*_2_), …)^⊤^ *and π*(*t*, 𝒳) ≜ (*π*(*t, x*_1_), *π*(*t, x*_2_), …)^⊤^. *Then, under the conditions* (2), (11), *and* (12), *Algorithm 1 almost surely has an estimation error*

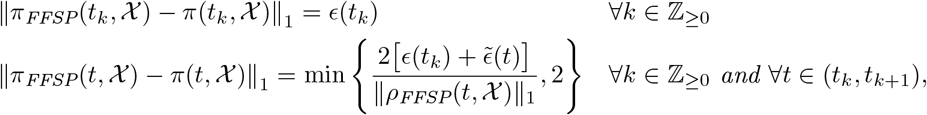

*where*

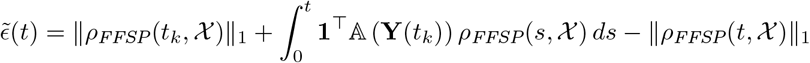

*for all k* ∈ ℤ_≥0_ *and t* ∈ (*t*_*k*_, *t*_*k*+1_), *and*

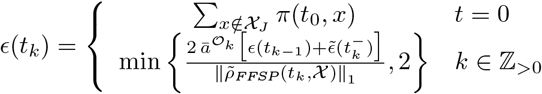

*with* 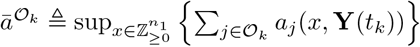.

The proof of this theorem is provided in the supporting material, specifically in section **S.2.2**. A significant advantage of this proof is that the computable error bound it offers relies solely on the FFSP solution. The error bound provided in Theorem 2 is computable because the FFSP solution *π*(*t*, 𝒳) contains only finitely many non-zero elements at each time point. This computable error bound allows us to evaluate the performance of the FFSP algorithm and assess its reliability. Moreover, as the size of the truncated state space 𝒳_*J*_ approaches the full (infinite) state-space 𝒳, both *ϵ*(*t*_0_) and 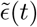 tend toward zero. This suggests that *ϵ*(*t*_*k*_) for *k* = 1, 2, …, also converges to zero, indicating that the accuracy of the algorithm can be indefinitely improved by expanding the size of the truncated state space. However, a major drawback of this error bound is its exponential growth in response to observable jump events, as *ϵ*(*t*_*k*_) is multiplied by 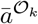 during error updating. This exponential growth can cause the error bound to reach large values quickly in some scenarios (see Section 3).

However, a significant challenge in this proof is analyzing the discrepancy between the un-normalized filter, *ρ*(*t, x*), and the FFSP estimate, *ρ*_FFSP_(*t, x*). Since the total mass of *ρ*(*t, x*) is not conserved, traditional techniques such as those in [15] are inapplicable. Instead, we propose an extended FSP theorem that leverages the structure of 𝔸(**Y**(*t*_*k*_)), notably the Metzler matrix property and the non-positivity of its column sums. This extended FSP theorem is crucial in proving Theorem 2 and, due to its versatile form, may be applicable in other scenarios where, for example, the initial distribution is uncertain or the total mass of the distribution is not conserved, as demonstrated in [52]. In [52], a non-linear partial differential equation is formulated to describe the probability of protein concentration in a population of dividing cells. Given the complexity of this non-linear equation, a linear version is proposed, featuring a non-constant total mass. The extended FSP theorem is thus well-suited to more accurately characterize the evolution of total mass in such contexts.

#### Theorem 3

(Extended FSP theorem for un-normalised probability distributions). *We consider a set of un-normalised probability distributions* 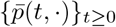 *defined on a discrete state space* 𝒳 = {*x*_1_, *x*_2_, …} *and evolving according to*

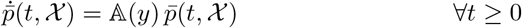

*where* 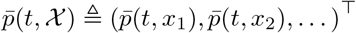, *the variable y is a vector in* 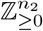, *and the matrix* 𝔸 (*y*) *is defined in* (9). *We also consider an FSP system of this infinite dimensional ODE, denoted by* {*p*_*FSP*_(*t*,·)} _*t*≥0_, *which is defined on the same state space but evolves according to*

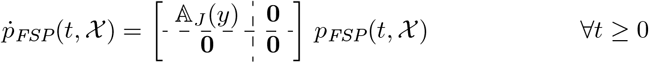

*where p*_*FSP*_(*t*, 𝒳) ≜ (*p*_*FSP*_(*t, x*_1_), *p*_*FSP*_(*t, x*_2_), …)^⊤^, *the matrix* 𝔸_*J*_ (*y*) *is the first J × J sub-matrix of* 𝔸 (*y*), *and p*_*FSP*_(0, *x*) = 0 *for all x* ∉ 𝒳_*J*_.

*Then, under conditions* (12) *and* 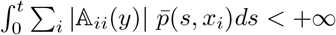, *the difference in the L*_1_ *norm between these two sets of measures can be evaluated by*

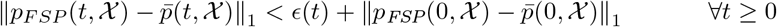

*where* 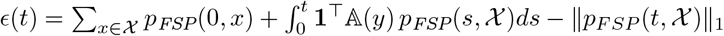. *(Here, ϵ*(*t*) *is computable because the function p*_*F SP*_ (*t*, ·) *has finite support*.) □

*Proof*. The proof is given in the section **S.2.1** of the supplementary material. Thanks to theorem Theorem 3, we were able to establish an error bound for Algorithm 1. However, as previously noted, the exponential growth of the error complicates assessing the actual accuracy of the algorithm. Although a large error bar does not necessarily indicate that Algorithm 1 is inaccurate—as evidenced by numerous examples in Section 3—it does tend to undermine confidence in this computational approach among practitioners. To address this issue, we next introduce an alternative FFSP algorithm that offers a tighter error bound, albeit with more limited applicability.

### 2.4 Another FFSP algorithm with tighter error bounds

In this section, we introduce a new FFSP algorithm designed to achieve tighter error bounds. In [15], the error of the original FSP algorithm can be precisely calculated due to the dominance property, where the FSP solution is consistently less than or equal to the exact solution component-wise. However, this property is lost in Algorithm 1 immediately after the normalization step. Once the total mass of the FFSP solution equals one, it cannot be universally dominated by any other probability distribution unless they are exactly identical, which is rare. This loss complicates the error analysis for Algorithm 1 and results in a somewhat conservative error bound as documented in Theorem 2.

The first FFSP algorithm utilizes a straightforward strategy ideal for addressing the filtering problem in noise-free scenarios and is versatile enough for most chemical reacting systems. Nevertheless, it yields a conservative error estimate for the approximate solution. To refine this, our new algorithm modifies the normalization step to ensure the exact solution always dominates the FFSP solution, allowing us to explicitly define the error. Specifically, when approximating the filter at the next observable jump time, this algorithm divides *ρ*_FFSP_(*t*_*k*+1_, *x*) by a quantity that exceeds the denominator used in (5) rather than the sum 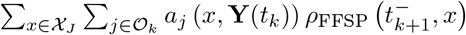. It is crucial to note that both algorithms yield the same result up to a normalization constant. While the first algorithm adopts the most intuitive approach for the noise-free filtering problem, the second employs a distinct normalization strategy that leads to an exact error certificate for the approximate solution. However, a limitation of the second algorithm is that the stochastic reaction network it applies to must meet specific conditions, as outlined in (11).

We present the new computational approach in Algorithm 2 and its estimation error in Theorem 4

#### Theorem 4.

*Under conditions* (2) *and* (11), *Algorithm 2 has the properties that*

- *π*_*FFSP*_(*t, x*) ≤ *π*(*t, x*) *for all t* ≥ 0 *and* 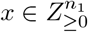, *and, therefore*,
- *the estimation error is given by* ∥*π*_*FFSP*_(*t*, 𝒳) − *π*(*t*, 𝒳)∥_1_ = 1 − ∥*π*_*FFSP*_(*t*, 𝒳_*J*_)∥_1_ *for all t* ≥ 0, *where π*_*FFSP*_(*t*, 𝒳) ≜ (*π*_*FFSP*_(*t, x*_1_), *π*_*FFSP*_(*t, x*_2_), …)^⊤^ *and π*(*t*, 𝒳) ≜ (*π*(*t, x*_1_), *π*(*t, x*_2_), …)^⊤^.

*Proof*. The proof is given in section **S.2.1**. □

The second part of Theorem 4 tells that we can exactly estimate the error of Algorithm 2 rather than obtaining a conservative error bound. This benefit mainly comes from the new normalization step, where we divide the un-normalised FFSP solution by a quantity larger than the actual normalization factors in (4) and (5). In addition to this theoretical result, our numerical examples in Section 3 also show that this second FFSP algorithm can provide a much tighter error bound than the first algorithm. All these results suggest that, the second FFSP algorithm (Algorithm 2) can provide estimates of lower error bounds than the first FFSP algorithm (Algorithm 1).

#### Algorithm 2 Another FFSP algorithm with a tighter error bound only valid under condition (11)

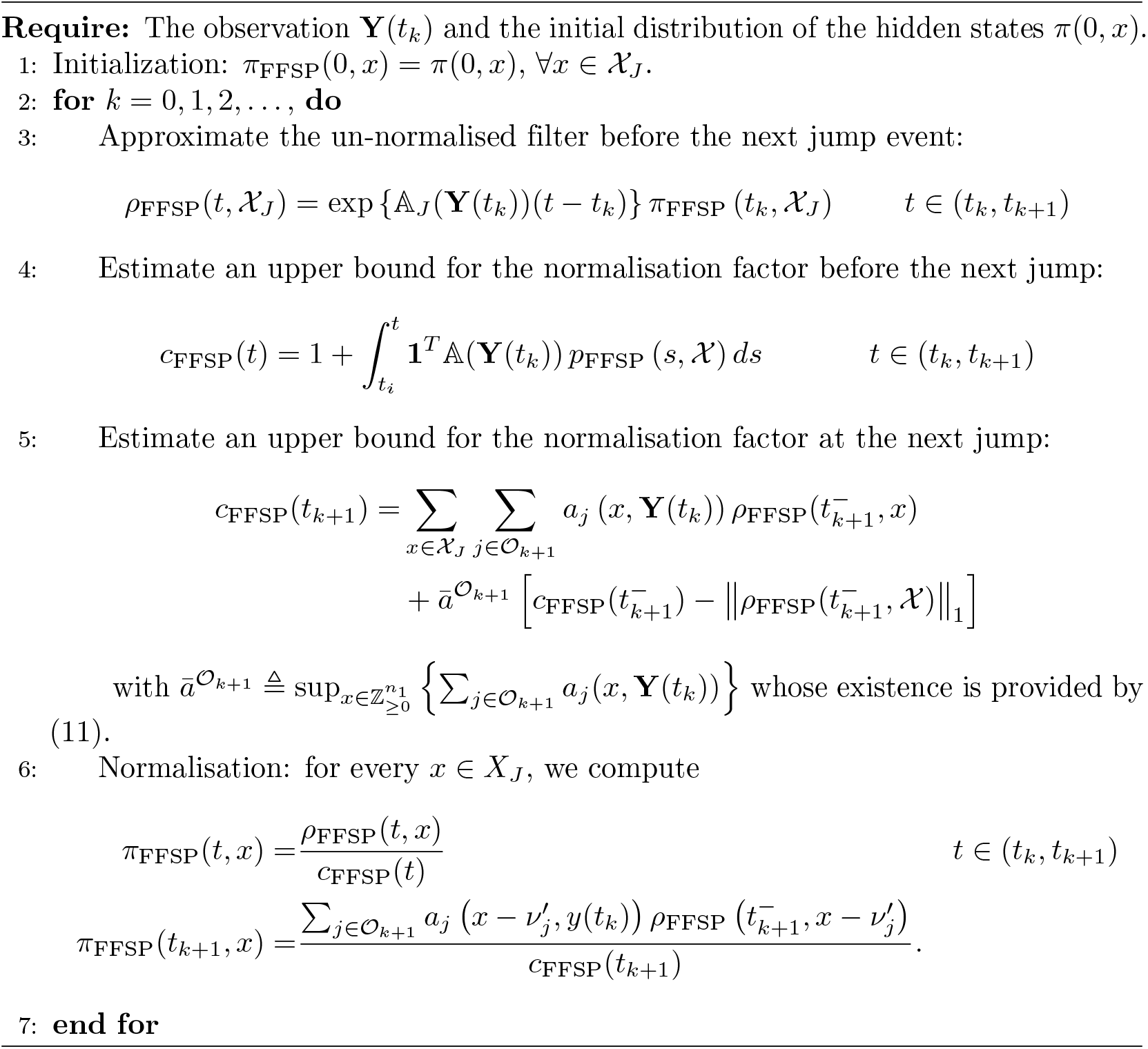

However, we need to mention that the tighter error bound of Algorithm 2 does not necessarily mean that this second FFSP algorithm is always more accurate than the first one. Mainly, this tighter error bound is obtained by a more elegant analysis technique which does not work for the first algorithm, but we cannot preclude the possibility that the first algorithm is actually more accurate than the second one. Our numerical examples in Section 3 also illustrate that the actual error of the first FFSP algorithm can be extremely tiny even though its error bound is very large. Moreover, by requiring the dominance property for the second FFSP algorithm, the solution of Algorithm 2 has a leaky property (similar to the original FSP in [15]) that its total mass constantly flows out of the truncated state space. Consequently, the second FFSP can lose most of its mass at some time point and stop being functioning after that. This leakiness is also observed in some numerical examples in Section 3, and in these cases, the first FFSP can perform much more accurately than the second one.

To summarise, Algorithm 1 and Algorithm 2 are equivalent up to a constant of normalisation, as their unnormalised filters follow the same linear equation and the algorithms only differs in the normalization step. The key difference is, that, the former employs a more natural strategy for solving the filtering problem, which gives rise to a probability distribution as an output. The latter, instead, utilises a different constant of normalisation to recover the dominance property, which therefore outputs an un-normalised probability distribution, which can then be normalised afterwards.

Also, the second FFSP has a narrower working range compared to the first FFSP algorithm. In the normalization step, the second FFSP algorithm requires the system to satisfy the condition (11), whereas the first FFSP algorithm does not. This suggests that the first FFSP can be applied to a broader class of chemical reacting systems. Nevertheless, both algorithms need condition (11) to generate error bounds.

Comparisons of these two FFSP algorithms are summarized in Table 1.

**Table 1.**
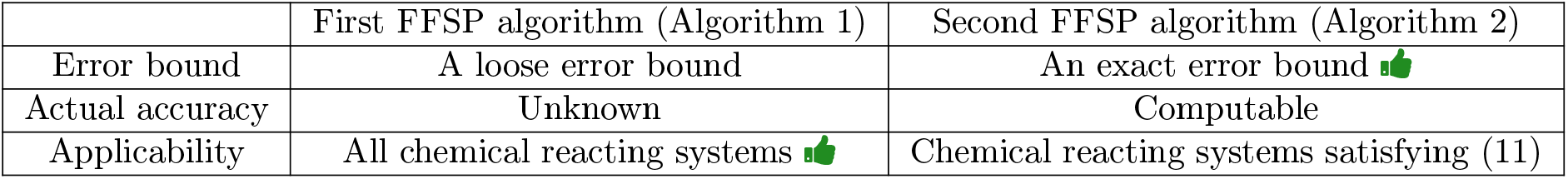
Comparisons of the two FFSP algorithm in terms of the error bound, actual accuracy, and applicability. The first FFSP algorithm can be applied to any chemical reacting system, while the second one requires the network to satisfy condition (11). However, both algorithms need condition (11) to be satisfied by the chemical reacting system in order to apply the accuracy analysis theorems.

## 3 Numerical Results

Now, we illustrate our filters using several biologically relevant examples. The experiments were performed on a laptop with a 2.3 GHz Dual-Core CPU, and the code is available on GitHub: Numerical Experiments Code.

### 3.1 A Simple Transcription-Translation Model

We start by considering a simple transcription-translation network containing basic patterns in the central dogma of molecular biology. The network consists of three reactions:

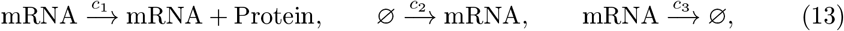

where an mRNA molecule translates proteins as well as being transcribed and degraded. We also assume that the protein degradation proceeds much slower than the above reactions so that in a relatively short time interval, the protein degradation can be ignored. In this example, we choose to continuously observe protein dynamics and estimate the mRNA copy number, and therefore we let **X**(*t*) ∈ ℤ_≥0_ keep track of the mRNA copy number and **Y**(*t*) ∈ ℤ_≥0_ keep track of the protein copy number.

#### 3.1.1 FFSP vs. Kalman Filter and Particle Filter

We first compare the performance of our FFSP to the classical Kalman filter and the particle filter introduced in [26]. We generated the hidden and observation process trajectories using a Gillespie simulation algorithm. To construct the Kalman filter, we approximate the dynamics of the concentrations 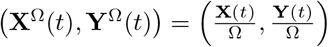 (where Ω is a sampling volume) with the diffusion approximation (see [19] for a review) and establish Kalman filters on top of it. Then, we used the generated observation process trajectory as input for the first FFSP algorithm (Algorithm 1) and the bootstrap particle filter (BPF); whereas, for the Kalman filter, we employed the same observation trajectory re-scaled with sampling volume Ω (see section **S3** of the Supporting Material for more details). The generated hidden process trajectory functions as a mean to test the quality of the estimates provided by the three filters. Moreover, we chose 𝒳_*J*_ = {0, 1, 500} as the truncated state space for the FFSP, *N*_*tot*_ = 10000 as the particle size of the particle filter, Ω = 100 as sampling volume, and *t*_*f*_ = 1*s* as final simulation time, and **Z**_0_ = [0, 0] as initial state. We assumed that the network follows mass-action kinetics and the propensity functions are *a*_1_(*z*_1_, *z*_2_) = *c*_1_*z*_1_, *a*_2_(*z*_1_, *z*_2_) = *c*_2_, *a*_3_(*z*_1_, *z*_2_) = *c*_3_*z*_1_. For the reaction rates, we set *c*_1_ = 100, *c*_2_ = 10, *c*_3_ = 5.

The numerical results are shown in Fig. 3. In this figure, the observation process is monotonically increasing as protein degradation is absent in our model. To evaluate the performance of the three filters, we plot all their estimates together with the exact trajectory of the hidden process.

**Fig. 3.**
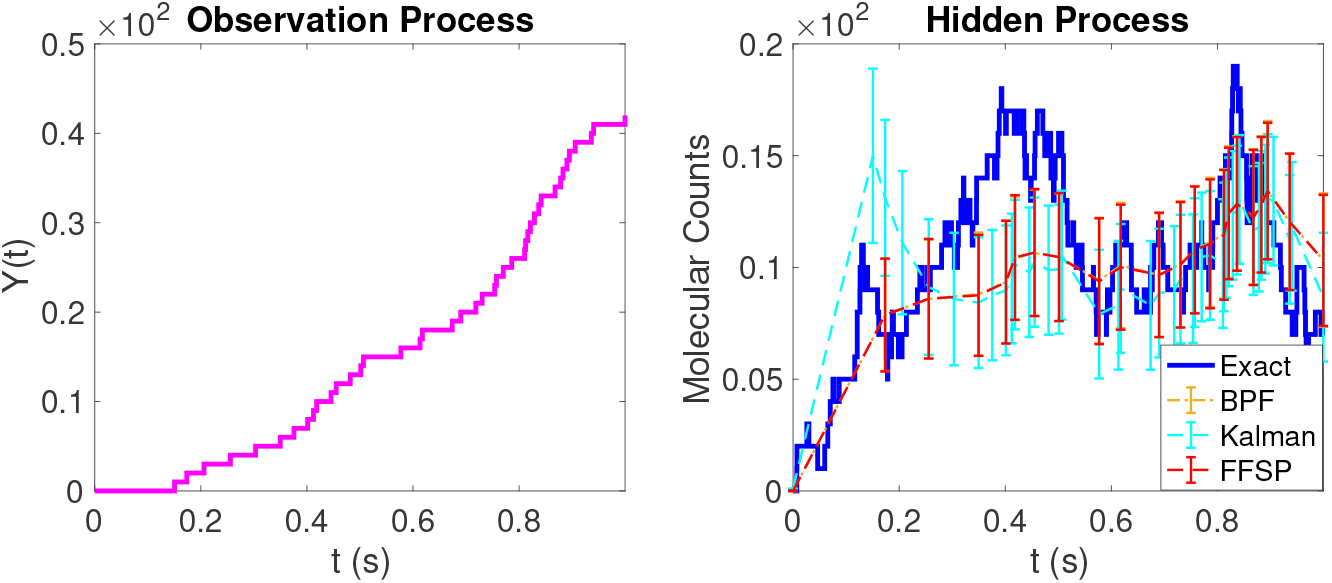
Performance of the FFSP, Kalman filter, and bootstrap particle filter (BPF) in the simple transcription-translation model. The first plot shows protein dynamics (the observation process) up to the final time. The second plot shows the exact trajectory of the mRNA dynamics (the hidden process) and the conditional mean estimates by three stochastic filters together with their standard deviations.

As shown, all filters capture the general trend of the hidden process well. Contrary to the introduction, the Kalman filter performs acceptably in reconstructing the trajectory of the hidden process for a linear network with no feedback. Additionally, the observation process trajectory resembles that of an SDE (Stochastic Differential Equation), given the moderately high copy number regimes, causing the filter to mistake Poisson-type noise for Gaussian-like noise. Therefore, in such scenarios, the Kalman filter remains a valuable inference tool. On the other hand, the FFSP and BPF show excellent agreement in their estimations.

#### 3.1.2 Comparison of two FFSP algorithms

We focus now on comparing the performance of the two proposed FFSP algorithms concerning the error propagation of the approximate filtering solution. To this end, we test the chemical reaction network in Eq. (13) with the same hidden (mRNA) and observed (protein) processes. To be consistent with the error analysis hypotheses, we chose the following propensity functions: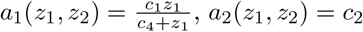, and *a*_3_(*z*_1_, *z*_2_) = *c*_3_*z*_1_ so that the propensity of the observable reaction (the first one) is upper bounded. The propensity functions of the observable reactions play a crucial role in the error bound behaviour (see Theorem 2). Therefore, for different values of the parameters *c*_1_ and *c*_4_, we may obtain distinct error bounds and dynamic scenarios. In Figs. 4 to 6 we show three different behaviours of the error bounds and of the dynamics over time. In all the three settings, we again simulated the observation and hidden process trajectories with a Gillespie algorithm and fed the former as input to both FFSPs algorithms. Furthermore, we set 𝒳_*J*_ = {0, 1, 500} as hidden species truncated state, **Z**_0_ = [5, 0] as initial condition and *t*_*f*_ = 2*s*.

**Fig. 4.**
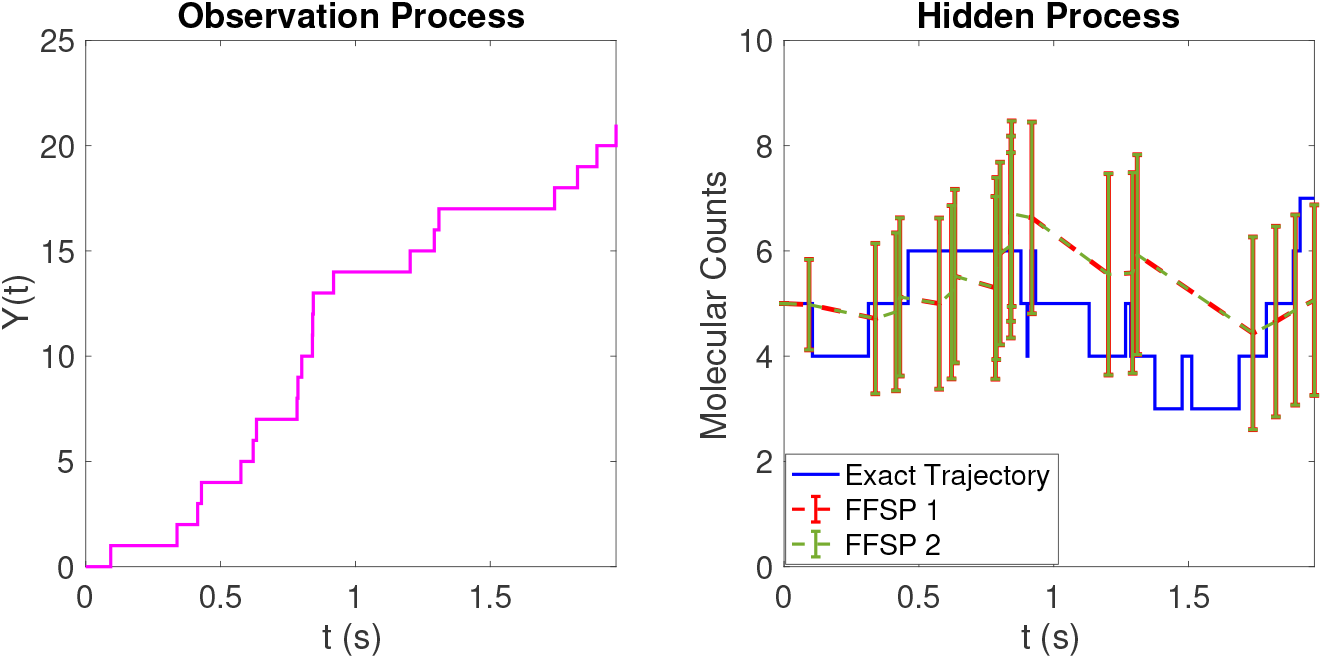
Comparison of the two FFSP algorithms in the simple transcription-translation model with a low protein production rate. The first plot shows the protein dynamics (the observation process) up to the final time. The second plot shows the exact trajectory of the mRNA dynamics (the hidden process) and the conditional mean estimates by the two FFSP algorithms (FFSP 1 and FFSP 2) together with their standard deviations. Specifically, the FFSP algorithms succeed in following the trend of the mRNA dyanmics and show a good agreement on their estimates, corroborating their equivalence.

### Scenario 1: Low protein production rate

In this first case, we set *c*_1_ = 25, *c*_2_ = 4, *c*_3_ = 1, and *c*_4_ = 10, so that the protein production rate saturates at a relatively low value 25. From the numerical result shown in Fig. 4, we can see that the observation process (the protein dynamics) still increases quite rapidly, in contrast to the mRNA copy number, which in turn fluctuates around the same value set as initial condition. Moreover, the two FFSP algorithms perform incredibly well in estimating the hidden process and show a very good agreement on all the estimates.

However, the error-bound growth tends to behave differently in the two algorithms (see Table 2).

**Table 2.** Error bounds of the two FFSP algorithms in case of the low protein production rate. The last line shows the *L*_1_ distance between the conditional distributions provided by the two FFSP algorithms.

| | $t_0$ | $t_1$ | $t_4$ | $t_8$ | $t_{14}$ | $t_{17}$ | $t_{21}$ |
| --- | --- | --- | --- | --- | --- | --- | --- |
| Jump times | 0 | 0.09 | 0.43 | 0.78 | 0.92 | 1.3 | 1.9 |
| 1<sup>st</sup> FFSP's error bound | 0 | $1 \times 10^{-1}$ | 2 | 2 | 2 | 2 | 2 |
| 2<sup>nd</sup> FFSP's error bound | 0 | $< 10^{-15}$ | $< 10^{-15}$ | $4.9 \times 10^{-15}$ | $8.2 \times 10^{-12}$ | $6.9 \times 10^{-9}$ | $1.6 \times 10^{-4}$ |
| Filter difference | 0 | $< 10^{-15}$ | $< 10^{-15}$ | $4.9 \times 10^{-15}$ | $8.2 \times 10^{-12}$ | $6.9 \times 10^{-9}$ | $1.6 \times 10^{-4}$ |

For this choice of parameters, the propensity *a*_1_(*z*_1_, *z*_2_) tends to have lower values at each jump time; therefore, every time it gets pre-multiplied with the error of the previous jump time (see Theorem 2), there is less accumulation of the error over time. Nevertheless, as it can be seen in Table 2, the error bound of the first FFSP algorithm grows immediately after the first few jump times. Such a feature can be expected, given the conservative upper bound. On the contrary, the error bound of the second FFSP algorithm is incredibly small for the first few jump times; then, we can notice a moderately rapid growth, but it stays pretty low until the last jump time with an order of magnitude of 10^*−*4^).

The last row of Table 2 shows the *L*_1_ difference in the values of the conditional distributions obtained with the two FFSP algorithms, which suggests that these two filters provide very similar results. To be more specific, the difference between the two filters is almost indistinguishable until *t*_14_ (order of magnitude from 10^*−*15^ to 10^*−*12^) and slowly grows towards the end of the simulation, remaining appreciably low (10^*−*4^). We recall that Algorithm 1 and Algorithm 2 are equivalent up to a constant of normalisation, which is very different in the two algorithms. The diverse normalisation strategy might be responsible for the difference in the filters values displayed towards the end of the simulation. To summarise, the second FFSP algorithm is notably very accurate, and the estimates provided by the first FFSP algorithm are considerably close to the result obtained by second one. Therefore, we can claim that Algorithm 1 is also very accurate, even though its conservative error bound grows incredibly fast from the beginning. Moreover, it is misleading to say that one algorithm is better than the other. Specifically, with the triangle inequality, it is possible to show that the *L*_1_ distance of the approximate solution provided by Algorithm 1 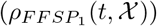 from the exact one *ρ*(*t, 𝒳*), namely 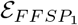, is included in the following interval 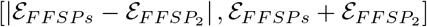. Here 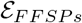 represents the *L*_1_ distance between the two approximate solutions obtained with Algorithm 1 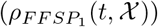 and Algorithm 2 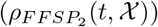 whereas, 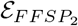 is the *L*_1_ distance between the approximate solution obtained with Algorithm 2 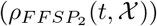 and the exact one *ρ*(*t, 𝒳*). Then, we can observe that, for example, during the early stages of the simulation (up to *t*_8_), 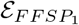 is included in an interval with a significant gap in the order of magnitude (10^*−*16^ to 10^*−*14^), while 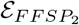 has the order of 10^*−*15^. Therefore, we may conclude that both filters are appreciably accurate.

### Scenario 2: High protein production rate

In this case, we set *c*_1_ = 40, *c*_2_ = 4, *c*_3_ = 1, *c*_4_ = 20 so that the protein production rate saturates at a relatively high value 40. In Fig. 5, we again show the observation process dynamics on the left side and the hidden dynamics, together with the filter estimates for the conditional means and standard deviations on the right. We can notice a large number of jump times in the protein dynamics, leading to rapid growth in a brief time. On the other hand, the mRNA copy number stays around the same value as the initial condition, like in the previous case. Again, the two FFSP algorithms show a good match in estimating the hidden process’ conditional expectations and standard deviations, but their estimates diverge after 1.8 seconds.

**Fig. 5.**
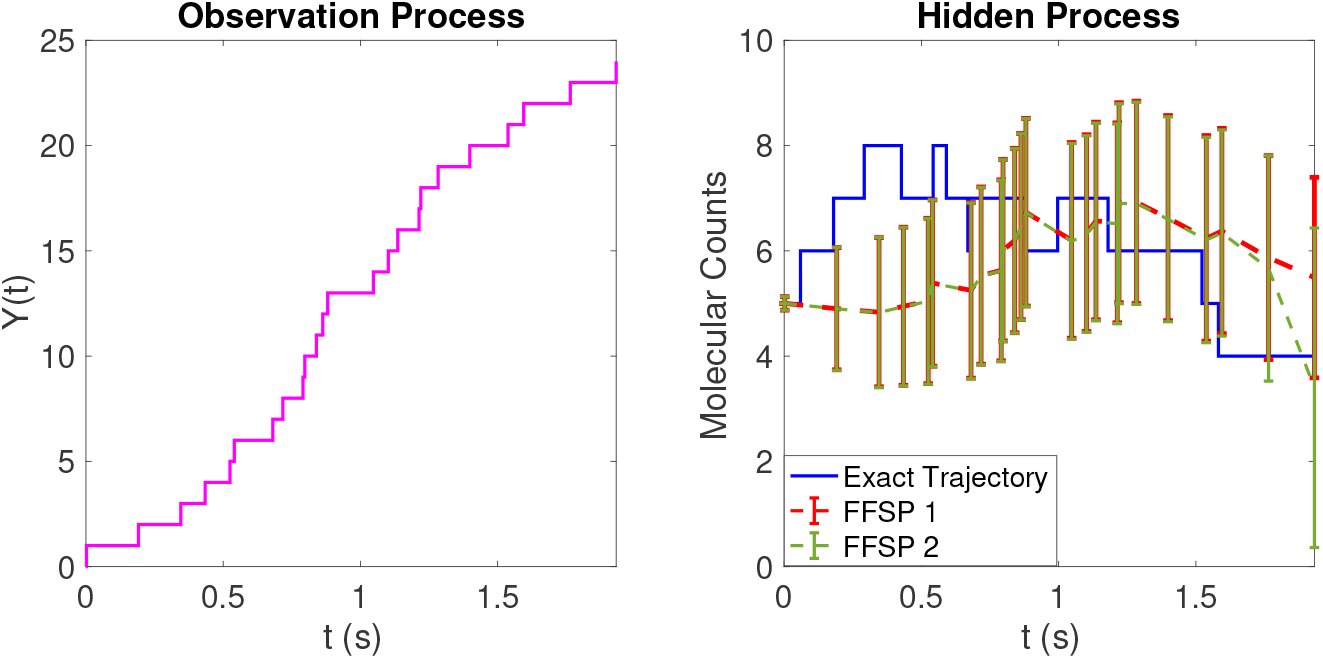
Comparison of the two FFSP algorithms in the simple transcription-translation model with a high protein production rate. The first plot shows protein dynamics (the observation process) up to the final time. The second plot shows the exact trajectory of the mRNA dynamics (the hidden process) and the conditional mean estimates by the two FFSP algorithm together with their standard deviations. Both the FFSP algorithms (FFSP 1 and FFSP 2) perform really well in estimating the hidden process dynamics, but their mean estimates diverge after around 1.8 seconds.

In this setting, the choice of parameters may lead to higher propensity values and, therefore, to a significant accumulation of the error bound over time. This is, in fact, confirmed by the error bound dynamics shown in Table 3. We can notice that the error bound of the first FFSP algorithm overgrows already from the start of the simulation. This behaviour is to be expected again, given the conservative error estimate. In contrast, the error bound of the second FFSP algorithm tends to be very low until the 18th jump time but then grows quite fast, reaching a relatively high order of magnitude (10^*−*1^). Even though the second FFSP algorithm is not as accurate as in the first case, it can still be considered accurate before 1.8 seconds. After 1.9 seconds, the second FFSP algorithm loses more than one third of the total mass and therefore can not be regarded as an accurate filter.

**Table 3.** Error bounds of the two FFSP algorithms in case of the high protein production rate. The last line shows the *L*_1_ distance between the conditional distributions provided by the two FFSP algorithms.

| | $t_0$ | $t_1$ | $t_2$ | $t_5$ | $t_{18}$ | $t_{21}$ | $t_{23}$ | $t_{24}$ |
| --- | --- | --- | --- | --- | --- | --- | --- | --- |
| Jump times | 0 | $1.8 \times 10^{-3}$ | 0.19 | 0.52 | 1.2 | 1.5 | 1.8 | 1.9 |
| 1<sup>st</sup> FFSP's error bound | 0 | $8.8 \times 10^{-5}$ | 0.47 | 2 | 2 | 2 | 2 | 2 |
| 2<sup>nd</sup> FFSP's error bound | 0 | $< 10^{-15}$ | $< 10^{-15}$ | $< 10^{-15}$ | $1.2 \times 10^{-7}$ | $1.9 \times 10^{-4}$ | $2.9 \times 10^{-2}$ | $3.7 \times 10^{-1}$ |
| Filters difference | 0 | $< 10^{-15}$ | $< 10^{-15}$ | $< 10^{-15}$ | $1.2 \times 10^{-7}$ | $1.9 \times 10^{-4}$ | $2.9 \times 10^{-2}$ | $3.8 \times 10^{-1}$ |

Despite this large gap in error bounds, the difference in the conditional distribution estimated values obtained by the two filters (last row of Table 3) again shows that the two algorithms produce appreciably similar approximated solutions up to *t*_18_, when the second filter is still accurate. Therefore, until *t*_18_, we can assert that both filters produce truly accurate estimates, without preferring one filter over the other, for the same reasons as in **Scenario 1**. Afterwards, the second filter tends to become inaccurate and we do not have means for discussing the accuracy of the first filter, given its poor error bound. In summary, we can conclude that the values of the observable propensity functions and the number of jumps of the observation process deeply impact the accuracy of the filters.

### Example 3: High protein production rate but less observable jumps

Now, we investigate how different observation trajectories affect the filters’ performance. To this end, we still keep *c*_1_ = 40, *c*_2_ = 4, *c*_3_ = 1, *c*_4_ = 20 as in the previous example but collect a simulation where the observation process jumps less frequently. In this case, we should observe less error propagation over time, and the error bounds should attain lower values at the final jump time.

As can be seen in Fig. 6, the two FFSP algorithms satisfactorily capture the behaviour of the hidden process. However, as shown in Table 4, the error bound of the first FFSP algorithm still grows very fast after the second jump time; whereas the error bound of the second FFSP algorithm behaves overall nicely and tends to grow slower than in the previous example, attaining an order of magnitude of 10^*−*4^ at the final time. From these results, we can conclude that the for the same system, the error bound of the second FFSP algorithm is largely influenced by the shape of the observation trajectory, especially the number of observable jumps. On the contrast, the error bound of the first FFSP algorithm does not show such an influence.

**Table 4.** Error bounds of the two FFSP algorithms in case of the high protein production rate but less observable jumps. The last line shows the *L*_1_ distance between the conditional distributions provided by the two FFSP algorithms.

| | $t_0$ | $t_1$ | $t_2$ | $t_3$ | $t_5$ | $t_8$ | $t_{10}$ | $t_{11}(=t_f)$ |
| --- | --- | --- | --- | --- | --- | --- | --- | --- |
| Jump times | 0 | 0.033 | 0.31 | 0.39 | 0.83 | 1.2 | 1.5 | 1.7 |
| 1<sup>st</sup> FFSP's error bound | 0 | $2.4 \times 10^{-2}$ | 2 | 2 | 2 | 2 | 2 | 2 |
| 2<sup>nd</sup> FFSP's error bound | 0 | $< 10^{-15}$ | $< 10^{-15}$ | $2.1 \times 10^{-15}$ | $1.5 \times 10^{-12}$ | $2.7 \times 10^{-9}$ | $8.6 \times 10^{-7}$ | $1.6 \times 10^{-4}$ |
| Filters difference | 0 | $< 10^{-15}$ | $< 10^{-15}$ | $2.1 \times 10^{-15}$ | $1.4 \times 10^{-12}$ | $2.7 \times 10^{-9}$ | $2.5 \times 10^{-5}$ | $1.6 \times 10^{-4}$ |

**Fig. 6.**
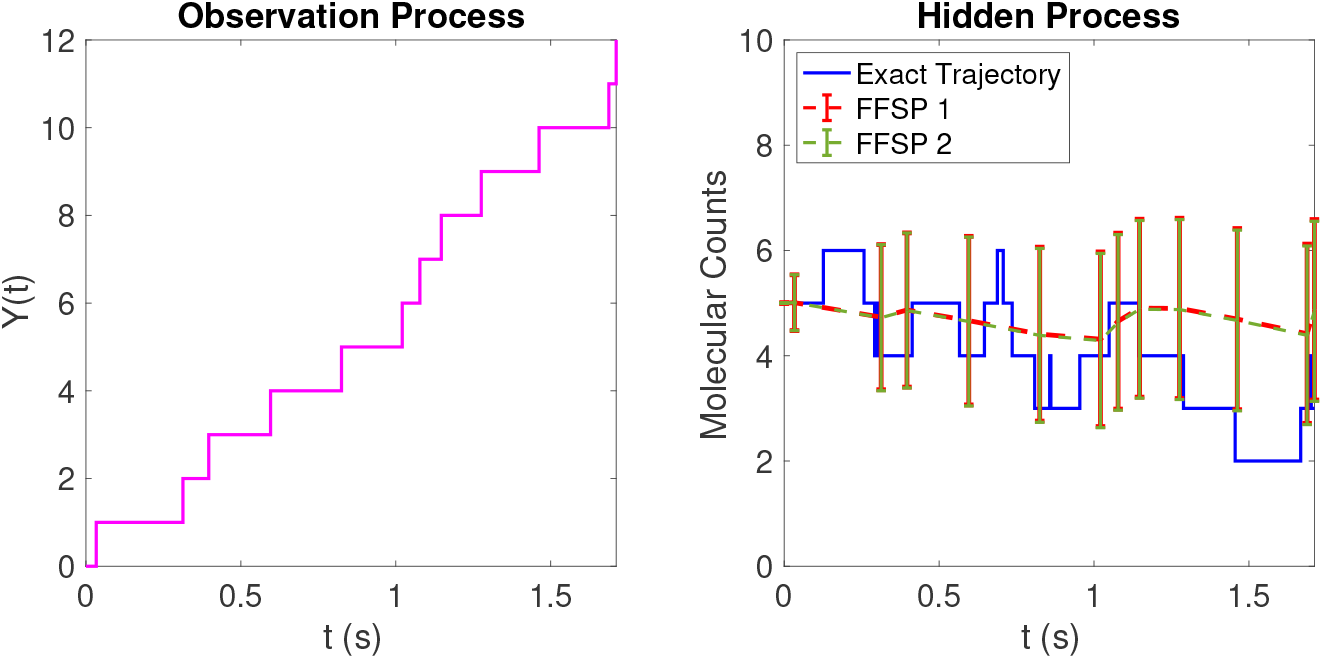
Comparison of the two FFSP algorithms in the simple transcription-translation model with a high protein production rate but less observable jumps. The first plot shows protein dynamics (the observation process) up to the final time. The second plot shows the exact trajectory of the mRNA dynamics (the hidden process) and the conditional mean estimates by the two stochastic filters together with their standard deviations. The two FFSP algorithms (FFSP 1 and FFSP 2) agree on all their estimates of the hidden process.

#### 3.1.3 FFSP vs. Particle filter: Conditional Distribution Estimation

Now, we consider the same network (13) but switch the observed and hidden species, i.e., we let **X**(*t*) keep track of the protein copy number and **Y**(*t*) of the mRNA copy number. In this setting, the protein copy number is solely determined by the first reaction channel, and therefore the conditional distribution *π*(*t, x*) follows a Poisson distribution with parameter 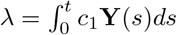. We again assume mass-action kinetics for the network and therefore the propensity functions remain the same as the first example. However, for reaction rates parameters, we set *c*_1_ = 1, *c*_2_ = 5, *c*_3_ = 1.

In this case, we test the performance of the FFSP filter (Algorithm 1) against the particle filter [26] in estimating the hidden dynamics, together with the conditional distribution. We again employed the Gillespie algorithm to generate the hidden and observed trajectories, and we then used the generated observation trajectory as input for both filters. We set the hidden species state space to be 𝒳 = {0, 1, 200} for the FFSP filter, *N*_*tot*_ = 10000 as particle size, *t*_*f*_ = 5*s* as final time and **Z**_0_ = [0, 5] as initial state. The numerical experiments are then shown in Fig. 7.

**Fig. 7.**
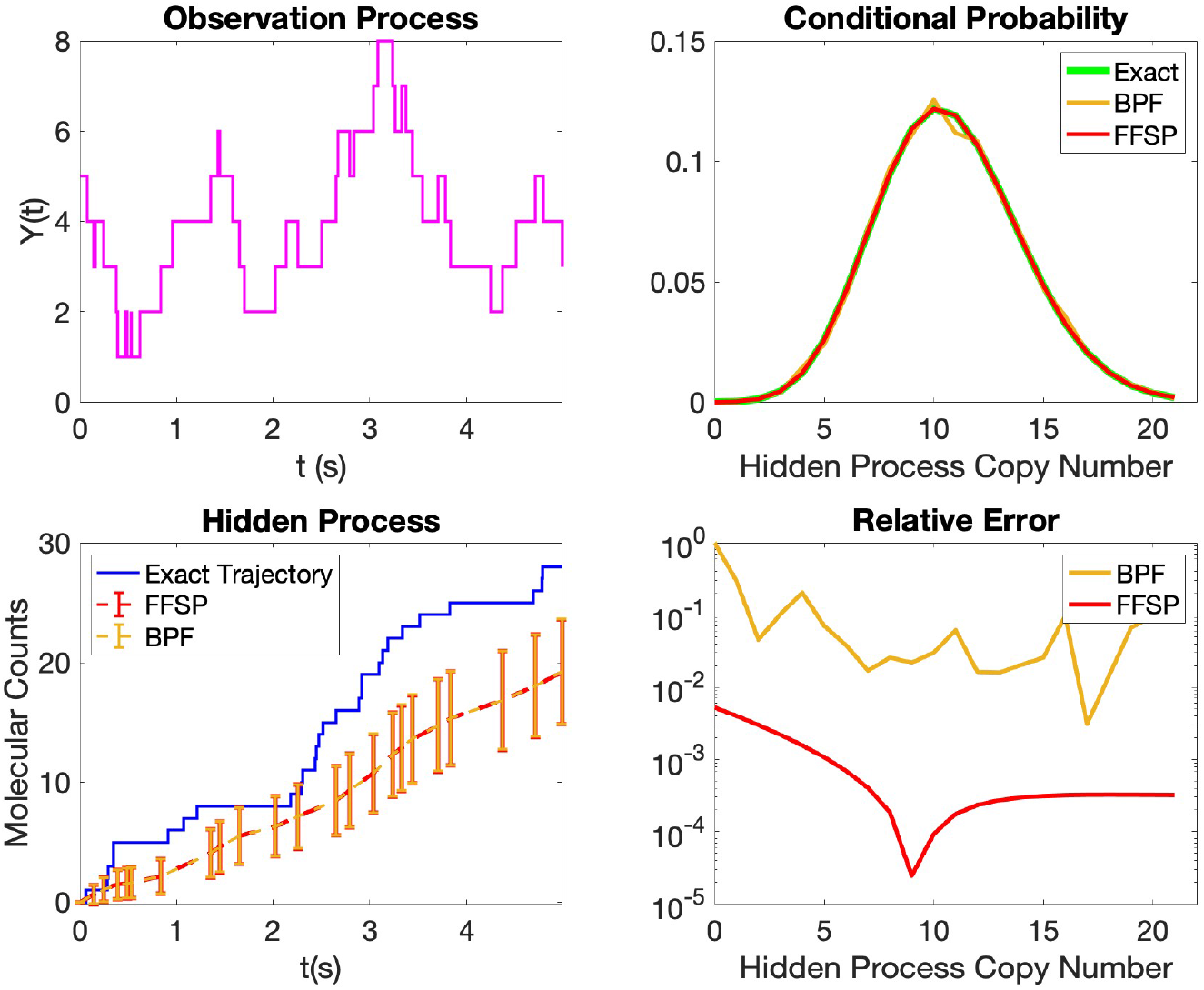
Comparison of the FFSP and bootstrap particle filter (BPF) in the simple transcription-translation model (observing the mRNA dynamics). The left column shows the observation and hidden process trajectories, together with the stochastic filters estimates and standard deviations, until the final simulation time. The top right plot, instead, displays a comparison between the exact conditional distribution and the ones estimated by the FFSP (Algorithm 1) and the particle filter at t=3s. The bottom right plot, instead, depicts the relative error of the filters from the exact conditional distribution. The result shows that the two filters have a similar performance in estimating the conditional mean and variance, but the FFSP is far more accurate in approximating the exact conditional distribution. Moreover, the particle filter takes 29.9s to be executed and the FFSP, in contrast, 11.3s, being 3 times faster.

As we can notice in the top left plot, the mRNA copy number fluctuates over time, with rapid outbursts that enhance protein translation. In this regard, we plotted the hidden process (protein) copy number in the bottom left plot as a validation of the filters estimates (conditional expectations). We can observe that the protein copy number increases rapidly over time, with the exception of some regions in which the mRNA copy numbers suddenly decreases. Overall, the two filters agree on all the estimates over time, with a general underestimation of the actual values towards at the beginning and at the end of the simulation analysis. The underestimation of the protein copy number might be due to the fact that the protein increases very rapidly at the initial time even though the mRNA copies tend to decrease. Therefore, the filters fail to follow this unexpected sharp increase at the beginning and keep underestimating the values for this whole interval.

Although the particle filter performs accurately in estimating the conditional means and standard deviations, we can discern, in the top right plot, that FFSP is more accurate in reproducing the exact conditional distribution, e.g., when estimating the peak probability. Such a behaviour can be expected, as the FFSP is a direct method, and therefore more accurate at depicting the behaviour of probability distributions. On the contrary, being particle filter a Monte-Carlo method, its performance depends on the number of particles being employed in the algorithm and how well they locate at a certain state. More specifically, let *V*_1_(*t*), …,*V*_*N*_(*t*) be *N* particles sampled from the conditional distribution; a particle filter use a scaled binomial distribution 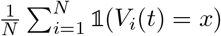 to approximate the conditional probability at the state *x*. In this case, the coefficient of error (the ratio between the standard deviation and the mean of this scaled binomial distribution) becomes 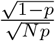 with *p* being the probability of finding the particle in that specific state, suggesting that the particle filter is not particularly suited in estimating rare events when *p* is small. This phenomenon is spotted in the bottom right plot of Fig. 7: the particle filter has very big relative error when estimating the state *x* = 0 whose conditional probability value is approximately 4.5 *×*10^*−*5^. In contrast, the relative error of the FFSP is always considerably small, even in estimating rare events.

In conclusion, though the FFSP and the particle filter have similar performance in estimating the conditional mean and variance, the FFSP has a much higher accuracy when estimating specific conditional probability, especially the rare events.

### 3.2 Genetic Toggle Switch

We consider now the following network, also known in biology as Toggle Switch:

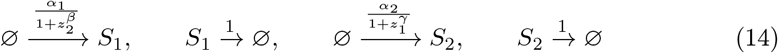

which is composed of two species mutually repressing each other. This elementary network shows properties like multi-stability and fluctuations that are desirable in different contexts in biology. Along these lines, a relatively new synthetic biology advance was implementing a genetic toggle switch in *Escherichia Coli* [53]. In this example, we decide to continuously observe the *S*_2_ copy number and estimate the dynamics of *S*_1_, and therefore we let **X**(*t*) ∈ ℤ _≥0_ keep track of the *S*_1_ copy number and **Y**(*t*) *∈* ℤ _*≥*0_ of *S*_2_. To be consistent with the mutual repression of the two species, we chose the subsequent propensity functions: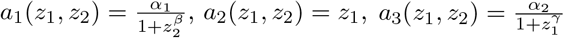, and *a*_4_(*z*_1_, *z*_2_) = *z*_2_.

The following scenarios test the two FFSP algorithms on the latent states’ estimates and error bound computations. Specifically, we chose two different parameter sets so that distinct dynamics and accuracy analyses arise. In all the examples, we set **Z**_0_ = [0, 0] as initial state and 𝒳 = {0, 200} as hidden species truncated state space and *t*_*f*_ = 5*s* as final simulation time. We again generated the observed and hidden process trajectories with a Gillespie algorithm and fed the former as input to both the filters.

#### Strong Repression

In this case, we set *α*_1_ = 16, *α*_2_ = 14, *β* = 1, *γ* = 1 so that the the two species strongly repress each other’s production. For this choice of parameters, switching behaviour is almost always guaranteed. Any time one of the two species is present in higher abundances, the other species’ values shrink to zero almost immediately. Such a behaviour can be seen in Fig. 8. The left-sided plot shows an initial slow increase of the observed (*S*_2_) species until time *t* = 4*s*, with a sharp outburst towards the end. In contrast, the right-sided plot shows the rapid increase of the hidden process from the beginning of the simulation and a rapid shrinking of its values towards the simulation’s final time, as expected. Moreover, the right-sided plot, in Fig. 8, also shows the stochastic filters’ estimates and their standard deviations. Generally, we can see that both FFSP filters capture the hidden process dynamics everywhere; for the error behaviour, both filters behave nicely, providing very low error bounds, especially second FFSP algorithm whose error is always below 10^*−*15^. In this case, we cannot distinguish whether it is an error of our algorithm or an error of the floating point.

**Fig. 8.**
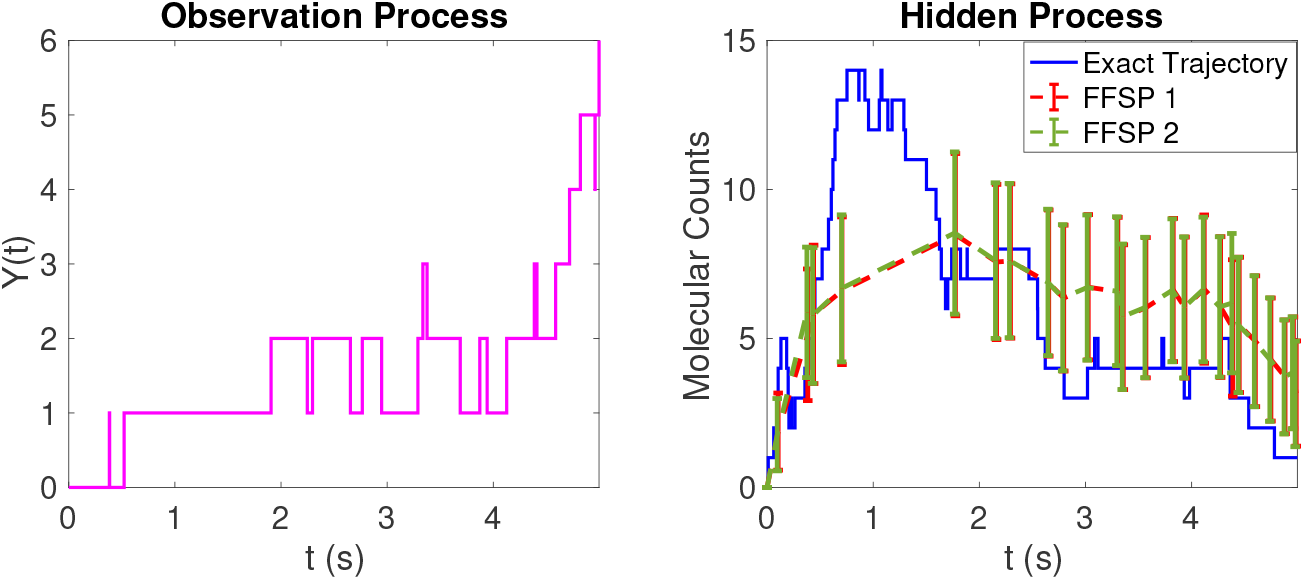
Performance of the two FFSP algorithms in the genetic toggle switch model (strong repression). The first plot shows the *S*_2_ (the observation process) copy number up to the final time. The second plot shows the exact trajectory of the *S*_1_ dynamics (the hidden process) and the conditional mean estimates by the two stochastic filters together with their standard deviations. The two FFSP algorithms (FFSP 1 and FFSP 2 in the figure’s legend) agree on all the estimations of the hidden process and succeed in following the trend of the hidden process.

#### Mild Repression

In this case, we chose *α*_1_ = 18, *α*_2_ = 15, *β* = 1.5, *γ* = 0.01, so that the first species repress the second one very mildly. In Fig. 9, we plot the observed and hidden process trajectories and filters’ estimates, together with their standard deviations (on the left and right side of the plot, respectively). In the first part of the simulation, both filters agree on the estimations and succeed in following the trend of the hidden process. However, both filters underestimate the outbreak of the hidden species because the size of this outbreak is unexpectedly large. Even when there is no repression, every second there are 7.5 newly produced *S*_2_ copies in average, which is only half of the size of this outbreak (see Fig. 9). In the second part of the simulation, when the hidden process values shrink, both filters can capture such a rapid decrease.

**Fig. 9.**
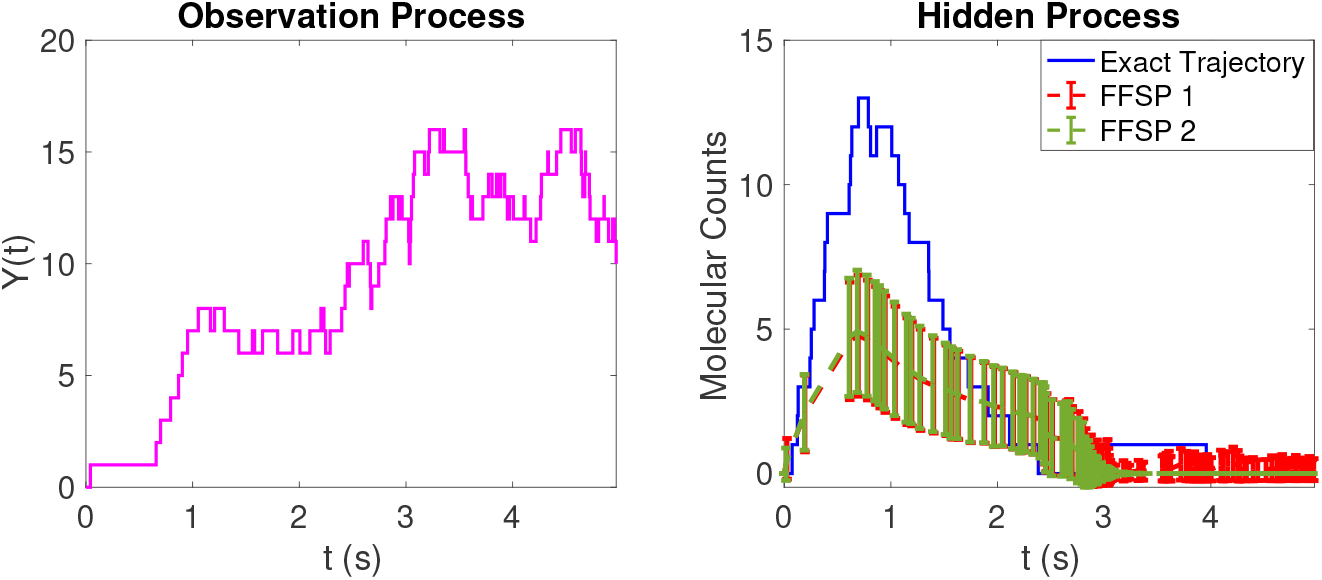
Performance of two FFSP algorithms in the genetic toggle switch model (mild repression). The first plot shows the *S*_2_ (the observation process) copy number up to the final time. The second plot shows the exact trajectory of the *S*_1_ dynamics (the hidden process) and the conditional mean estimates by the two stochastic filters together with their standard deviations. The two FFSP algorithms (FFSP 1 and FFSP 2 in the figure’s legend) agree on all the estimations of the hidden process until time *t* = 3*s*, after which only the first FFSP filter is still able to better capture the latent states.

Initially, the two filters can be both very accurate: as it can be seen in Table 5, the difference in their estimates is incredibly small. Moreover, the second filter error bound tends to stay appreciably low until *t*_12_. Therefore, during the beginning of the simulation, both filters are to be considered very accurate. However, after 3 seconds, the second FFSP algorithm stops functioning as it loses almost all of its mass (see Fig. 9 and Table 5). In contrast, the first filter is functioning till the end though its error bound is very large (see Fig. 9 and Table 5). To conclude, there is no strict preference among the two filters during the early stages of the simulation; the first FFSP algorithm is preferred over the final time, when it can still capture the behaviour of the hidden process despite the large error bound dynamics.

**Table 5.** Error behaviour of the two FFSP algorithms in the example of genetic toggle switch with mild inhibition.

| | $t_0$ | $t_1$ | $t_2$ | $t_5$ | $t_{12}$ | $t_{21}$ | $t_{42}$ | $t_{60}$ |
| --- | --- | --- | --- | --- | --- | --- | --- | --- |
| Jump times | 0 | $4.1 \times 10^{-2}$ | $6.6 \times 10^{-1}$ | $8.7 \times 10^{-1}$ | 1.4 | 2.2 | 2.9 | 3.6 |
| 1<sup>st</sup> FFSP's error bound | 0 | $2.6 \times 10^{-2}$ | $9.6 \times 10^{-1}$ | 2 | 2 | 2 | 2 | 2 |
| 2<sup>nd</sup> FFSP's error bound | 0 | $< 10^{-15}$ | $3.6 \times 10^{-14}$ | $3.2 \times 10^{-13}$ | $1.2 \times 10^{-9}$ | $9 \times 10^{-5}$ | $7.8 \times 10^{-1}$ | 1 |
| Filters difference | 0 | $< 10^{-15}$ | $3.6 \times 10^{-14}$ | $3.2 \times 10^{-13}$ | $1.2 \times 10^{-9}$ | $9 \times 10^{-5}$ | $7.8 \times 10^{-1}$ | 1 |

### 3.3 Genetic Switch with Feedback: FFSP vs Particle Filter

We consider now the following network:

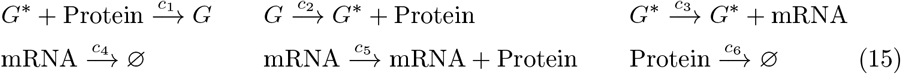

where *G* and *G*^*∗*^ are inactive and active genes, respectively activated and inactivated by the protein. Moreover, the active gene *G*^*∗*^ transcribes mRNA copy number, which can degrade. In turn, mRNA is responsible for the protein translation, which also degrades. For this network, we decided to employ mass-action kinetics and, thus, the propensity functions are *a*_1_(*z*_1_, *z*_2_, *z*_3_, *z*_4_) = *c*_1_*z*_2_*z*_4_, *a*_2_(*z*_1_, *z*_2_, *z*_3_, *z*_4_) = *c*_2_*z*_1_, *a*_3_(*z*_1_, *z*_2_, *z*_3_, *z*_4_) = *c*_3_*z*_2_, *a*_4_(*z*_1_, *z*_2_, *z*_3_, *z*_4_) = *c*_4_*z*_3_, *a*_5_(*z*_1_, *z*_2_, *z*_3_, *z*_4_) = *c*_5_*z*_3_, and *a*_6_(*z*_1_, *z*_2_, *z*_3_, *z*_4_) = *c*_4_*z*_3_. We continuously observe protein dynamics and estimate the other remaining species; therefore, we let 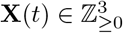 keep track of *G, G*^*∗*^, and the mRNA and **Y**(*t*) *∈* ℤ_*≥*0_ keep track of the protein.

In this numerical experiment, we compare the performance of the FFSP and the particle filter in estimating the various hidden species. To this end, we simulated the observed and hidden process trajectories with a Gillespie algorithm, and we fed the former as input to both filters. Moreover, we set *c*_1_ = 1, *c*_2_ = 5, *c*_3_ = 8, *c*_4_ = 2, *c*_5_ = 4, *c*_6_ = 1, *t*_*f*_ = 5*s* as final simulation time, and 𝒳 = {0, 100} as hidden species truncated state space for the mRNA copy number.

In Fig. 10, we can spot the observation and hidden process trajectories, together with the filters’ estimates and standard deviations. We can notice how the protein copy number starts with a slow increase, given that the active gene alternates periods of inactivity to activity and then experiences a sharp outburst once the gene stays mostly active. The same behaviour is overall held also by the mRNA copy number. Regarding the estimations carried out by the filters, we can see that the both filters perform well in tracking both the activated gene and mRNA dynamics.

**Fig. 10.**
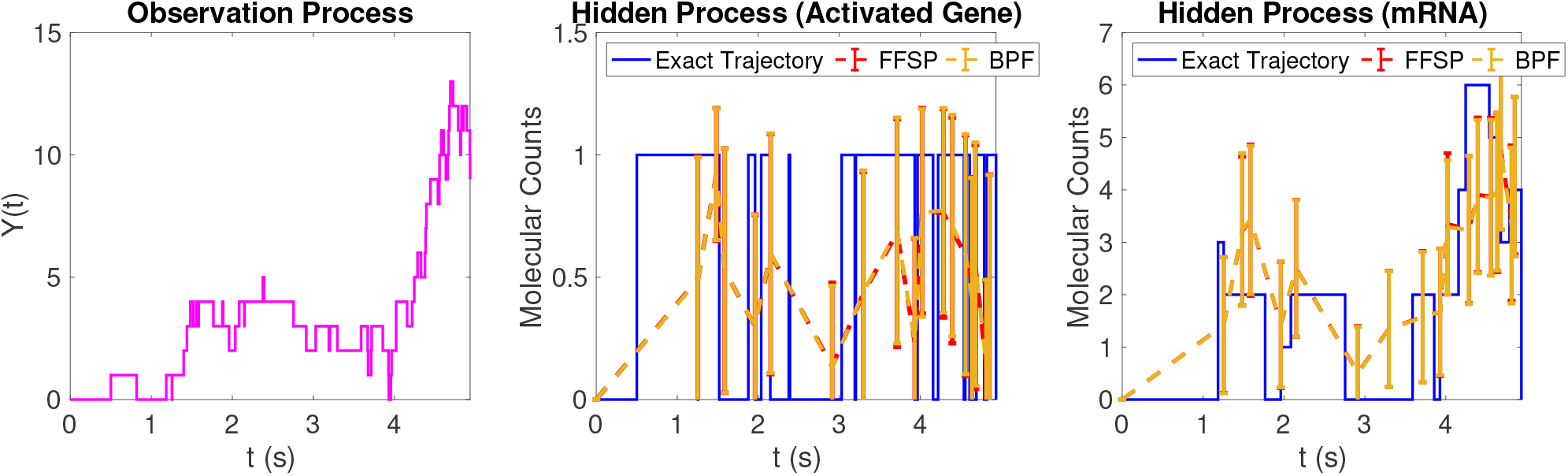
Performance of the FFSP and bootstrap particle filter (BPF) algorithms in the genetic switch model. The first plot shows the protein copy number up to the final time. The second and third plots show the exact trajectory of the activated gene and mRNA dynamics, respectively; also, we present their conditional mean estimates by the two stochastic filters together with the standard deviations. The FFSP shows a better performance in estimating both the activated gene and mRNA copy number over the particle filter, which instead underestimates the actual values overall.

### 3.4 Genetic Switch in a Hybrid Experimental Setup

In this subsection, we test the performance of the first FFSP algorithm in genetic switches operated in a hybrid experimental setup, where the protein dynamics is simulated in computers according to the trajectory of fluorescent mRNA molecules in real cells. With this aim, we consider the same genetic switch as before, but without the protein feedback:

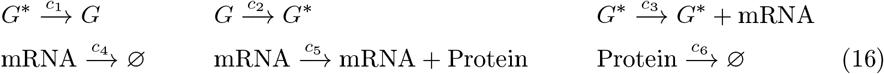

where *G* and *G*^*∗*^ are inactive and active genes. The active gene *G*^*∗*^ transcribes mRNA copy number, which can degrade. In turn, the mRNA is responsible for the protein translation, which also degrades. Given the lack of feedback, the gene switching together with the mRNA transcription parts of the circuit can be decoupled from the protein translation, resulting in the following reduced models:

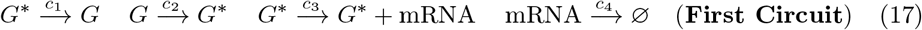

and

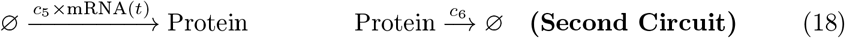

In particular, the first circuit (17) was implemented in yeast cells, which were made light-sensitive by tagging the mRNA molecules with fluorescent reporters. Upon a constant light stimulation, the mRNAs’ fluorescence was recorded for four hours under a microscope in an optogenetic platform [48, 49]. In the experiment, we collected data from 60 yeast cells simultaneously. The second circuit (18) was simulated *in-silico* through time-varying Gillespie algorithms. Mass-action kinetics were employed for the protein translation and degradation propensities. Furthermore, the mRNA trajectories from the circuit (17), acquired with microscope measurements, were used as inputs of distinct Gillespie algorithms to generate the protein trajectories. To be more specific, between the mRNA jump times *t*_*i*_ *≤ t < t*_*i*+1_, the mRNA molecular abundance, mRNA(*t*_*i*_), was fed as input parameter of the propensity function responsible for protein translation. Consequently the full protein trajectory was generated as a succession of iterative Gillespie algorithms with the different mRNA molecular abundances as inputs.

The main objective, then, is in using these protein trajectories as observation processes and predicting the mRNA dynamics with the first FFSP algorithm. To this end, we considered the whole circuit (16) and continuously observed protein dynamics and estimated the other remaining species; therefore, we let 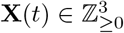 keep track of *G*^*∗*^, *G*, and the mRNA and **Y**(*t*) *∈* ℤ_*≥*0_ keep track of the protein. Mass action kinetics type propensities were used with the following parameters: *c*_1_ = 0.2, *c*_2_ = 0.1, *c*_3_ = 23, *c*_4_ = 0.5, *c*_5_ = 1, *c*_6_ = 1. Then, the initial condition was set to **X**_0_ = [0, 1, 0, 0]. The results are shown in Fig. 11. In particular, the top part shows the protein trajectory, generated with the Gillespie algorithm, with the corresponding mRNA abundance as an input; the bottom subplots display the *in-vivo* mRNA trajectories, together with the filters’ estimations. As can be seen in all the three representative cells, the first FFSP algorithm is able to reconstruct the *in-vivo* mRNA trajectories with a very good agreement, showing the potential use of the algorithm for real biological applications.

**Fig. 11.**
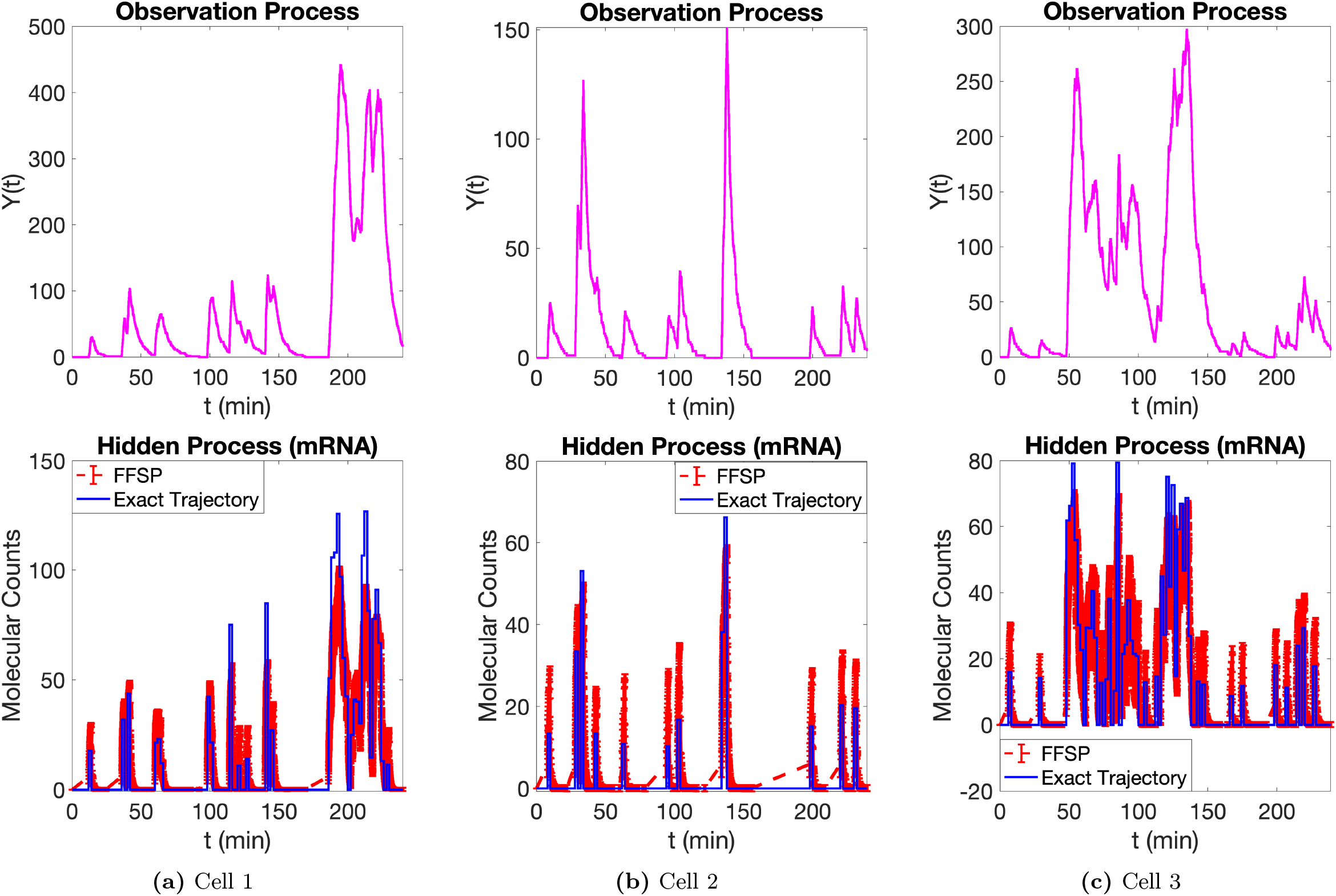
Performance of the FFSP algorithm in reconstructing the *in-vivo* mRNA trajectories of genetic switches operated in a hybrid experimental setup. The top subplots depict the protein trajectories employed as observation processes to predict the *in-vivo* mRNA trajectories. The bottom subplots show the in-vivo mRNA trajectories recorded with the microscope together with the FFSP estimates. The filter reproduces remarkably well all the three cases. Source data are provided in the source data file.

## 4 Conclusion

In this paper, we developed a direct approach to solve the filtering equation for a noise-free observation process in stochastic reaction networks. Specifically, we considered a continuous-time discrete-state Markov Chain 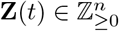 and assumed to exactly observe the dynamics of some chemical species. To postulate the filtering problem, we subsequently decomposed the network into 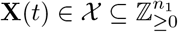 hidden species and 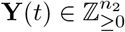, with *n*_2_ = *n – n*_1_, observed species. Thus, we assumed to exactly (noise-free) and time-continuously observe the trajectory of **Y**(*t*) and estimate the hidden species copy number **X**(*t*) dynamics. To be more specific, we wish to compute the conditional distribution *π*_*t*_ of the hidden process **X**(*t*) given the trajectory of the observed process **Y**(*s*) 0 *≤ s≤ t*. In this setting, such conditional distribution is a piece-wise deterministic Markov process (PDMP): it evolves according to a non-linear dynamical equation and jumps when the observation process does so.

This conditional distribution admits an un-normalised version that is also a PDMP and evolves according to a linear evolution equation between the observation process jump times. Moreover, the evolution matrix is similar to the generator matrix of the Chemical Master Equation (CME) and shares several valuable properties. Therefore, we solved the filtering problem by iteratively applying the Finite State Projection Method [15] to the linear system satisfied by the un-normalised distribution between the observation process jump times. We then recovered the filtering equation’s solution by normalising it before each jump time. This procedure gives rise to the Filtered Finite State Projection Method. Compared to the standard Finite State Projection method, the FFSP’s solution does not have a conserved total mass in time. To be able to carry out the accuracy analysis, we developed a modified FSP theorem to include also such distributions. The modified FSP theorem provides an exact error certificate for setting in which the initial probability distribution is not fully exact. Thanks to this theorem, we then then performed an accuracy analysis which results in a conservative error bound for the approximated solution. To resolve this issue, we provided an alternative algorithm, which exploits a different normalisation technique and results in an exact error certificate as in the FSP theorem [15], at a cost of narrower applicability.

From the numerical experiments in Section Section 3, we observe that the FFSP method accurately estimates the hidden dynamics of several biological networks. Moreover, in the introduction, we demonstrated via a numerical example that our method outperforms established filters, such as the Kalman and Extended Kalman filters, in complex nonlinear networks with feedback and Poisson-type noise. However, in the presence of a linear network and in higher copy number regimes where the observation process mimics Gaussian-type noise, the Kalman filter in section Section 3.1.1 shows performance similar to that of the FFSP and particle filters.

Additionally, the FFSP exhibits superior performance in estimating the full conditional distribution and rare events compared to the bootstrap particle filters, as shown in section Section 3.1.3. Furthermore, our filter produces accurate estimates when examining the error behavior of the proposed second algorithm. Finally, the FFSP algorithm performs well in reconstructing several *in vivo* mRNA trajectories recorded with a microscope in yeast cells for a genetic switch-type circuit, as seen in Section Section 3.4.

All these results suggest that our algorithm could be further employed to explore other biological circuits of interest, whose behaviors are still not fully understood in terms of latent state dynamics.

In the future, this work could be extended in several directions. One possible direction is to mitigate the curse of dimensionality. Since our algorithm has to solve an extensive linear system between each observation process jump time, the computational cost grows exponentially with the number of species. Therefore, developing more scalable approaches for high-dimensional filtering equation in the stochastic reaction network framework is needed. A possible approach may reside in using deep learning, as it has been shown to cope quite well with high-dimensional problems in the stochastic reaction networks framework [54] and other fields [55]. Second, the algorithm could further be extended by including parameters’ uncertainty, as the exact parameter values are usually unknown to scientists. Moreover, the FFSP can also be applied to solving CMEs with the Rao-Blackwell technique, which entails uncoupling the entire network into two parts and then marginalising out one by computing its conditional probability (see [56, 57] for the Rao-Blackwell technique and [45, 46] for the application of this idea together with moment closure to CMEs). The advantage of the Rao-Blackwell method is guaranteed by the Rao-Blackwell theorem (a variant of the law of total variance), suggesting that this method is more accurate than the Monte Carlo one under the same sample size. In this framework, our FFSP is ideal for the marginalisation part of the Rao-Blackwell technique when applied to biochemical reacting systems; some attempts in this direction were reported in [24].

## Supporting information

https://github.com/EleSofi/FFSP_2024_code_data

## Supporting information

**S1 Supplementary Material. The proofs of the theorems**. In the supplementary material, we rigorously verify the filtering equation under the assumed conditions and perform the error analysis for both FFSP algorithms.

**S2 Supplementary Material**. In this section of the supplementary material, we concern ourselves with the proofs of the error analysis results of Algorithm 1 and Algorithm 2.

**S3 Supplementary Material**. In this section, a derivation of the Kalman and Extended Kalman filters used in the introduction and in the numerical experiment 3.1.1 is shown.

## Acknowledgments

This work was funded by a grant from the Swiss National Science Foundation (Grant No. 182653).

## Footnotes

1 The well-posedness of *π*(*t, x*) is guaranteed by [29, Theorem 2.24], and we provide a more detailed discussion about it in the supplementary material.

