## Supplementary material for "Filtered finite state projection method for the analysis and estimation of stochastic biochemical reaction networks": https://github.com/EleSofi/FFSP_2024_code_data

#### S.1 Verification of the filtering equation

In this section, we prove the first theorem in our paper that under the non-explosivity condition

$$\sup_{t\in[0,T]} \sum_{j=1}^{M} \mathbb{E}\left[a_j^2(\mathbf{Z}(t))\right] < \infty \qquad \forall T > 0, \tag{S1}$$

the conditional distribution $\pi(t,x)$ uniquely solves the filtering equation up to indistinguishability. When the state space is finite (e.g., the chemical reaction system admits positive conservation laws), the result has already been shown in [1]; however, the result in the case of infinite state spaces was still open before this work. In Section S.1.1, we first discuss the well-posedness of the conditional distribution $\pi(t,x)$. Then in Section S.1.2, we rewrite the filtering equation into a form that is easier to analyze. Finally, we prove the well-posedness of the filtering equation in Section S.1.3 using the innovation method [2] and a Picard iteration [3, 4].

##### S.1.1 Well-posedness of $\pi(t,x)$

In this paper, we are interested in calculating the conditional distribution of hidden state $X(t)$ given the observations up to time $t$, i.e., $P\{\mathbf{X}(t) = x|\mathbf{Y}(s), 0 \le s \le t\}$. Here, we need to emphasize a few points. First, for every fixed $t$, this conditional distribution is random (rather than deterministic) and is measurable with respect to the filtration generated by $\mathbf{Y}(t)$, which we denote by $\mathcal{Y}_t$. In other words, the value of this conditional distribution fully depends on the random trajectory of the observation process. Second, for every fixed $t$, this conditional distribution is not uniquely defined. Mathematically, any $\mathcal{Y}_t$-adaptive random distribution $\mathcal{P}(x)$ can be viewed as a version of this conditional distribution as long as it satisfies

$$\mathbb{E}\left[\mathcal{P}(x)A\right] = \mathbb{E}\left[\mathbb{1}(\mathbf{X}(t) = x)A\right] \quad \forall \mathcal{Y}_t\text{-measurable random variable A with finite expectation and } \forall x \in \mathbb{Z}_{\ge 0}^{n_1}.$$

Consequently, $P\{\mathbf{X}(t) = x|\mathbf{Y}(s), 0 \le s \le t\}$ viewed as a continuous-time process is not uniquely defined either.

Among all these choices of conditional distributions, there is a càdlàg version, denoted by $\pi(t,x)$, satisfying $\lim_{s\to t^+} \pi(s,x) = \pi(t,x)$ and permitting the existence of $\lim_{s\to t^-} \pi(s,x)$ [2, Theorem 2.24]. In this paper, we specifically investigate such a conditional distribution. Moreover, the non-explosivity condition (S1) implies that the propensities $a_j(\mathbf{Z}(t))$ (for each $j = 1, \dots, M$) are uniformly integrable in any finite time interval $[0,T]$. Therefore, in addition to the càdlàg property of $\pi(t,x)$, the conditional expectation $\pi(t,a_j) \triangleq \sum_{x'} a_j(x', \mathbf{Y}(t))\pi(t,x')$ (for each $j = 1, \dots M$) is also càdlàg under the non-explosiviy condition (S1) [2, Remark 2.27].

##### S.1.2 Alternative form of the filtering equation

Here, we rewrite the filtering equation in a form that is easier to analyze. We recall that $\nu_i''$, $i \in \mathcal{O}$, are the sub-stoichiometry vectors regulating the net change of the observation process $\mathbf{Y}(t)$ once the reaction $i$ has fired. Such sub-vectors may not be all distinct. Therefore, in addition to the notations introduced in the main text, we further term

- $\{\mu_1, \ldots, \mu_{m_1}\}$ as the set of non-zero and distinguishable $\nu_i''$, $i \in \mathcal{O}$
- $\mathcal{O}_{\mu_k} \triangleq \{j | \nu_j'' = \mu_k\}$ $(k = 1, \ldots, m_1)$ as the set in which non-zero $\nu_j''$, $j \in \mathcal{O}$, are identical to $\mu_k$,
- $\tilde{R}_{\mu_k}(t) \triangleq \sum_{j \in \mathcal{O}_{\mu_k}} R_j \left( \int_0^t a_j(\mathbf{X}(s), \mathbf{Y}(s))\, ds \right)$ as the total firing number of the reactions in $\mathcal{O}_{\mu_k}$ up to time $t$,
- $a^{\mathcal{O}_{\mu_k}}\left(\mathbf{X}(s), \mathbf{Y}(s)\right) \triangleq \sum_{j \in \mathcal{O}_{\mu_k}} a_j\left(\mathbf{X}(s), \mathbf{Y}(s)\right)$ as the rate of the process $\tilde{R}_{\mu_k}(t)$.
- $a^{\mathcal{O}}\left(\mathbf{X}(s), \mathbf{Y}(s)\right) \triangleq \sum_{j \in \mathcal{O}} a_j\left(\mathbf{X}(s), \mathbf{Y}(s)\right)$ as the sum of the propensities of the observable reactions.

Then, the filtering equation can be rewritten as

$$\begin{aligned}
\pi(t,x) =& \pi(0,x) + \int_0^t \sum_{j \in \mathcal{U}} a_j(x - \nu_j', \mathbf{Y}(s)) \pi(s, x - \nu_j') - \sum_{j \in \mathcal{U}} a_j(x, \mathbf{Y}(s)) \pi(s,x) ds \\
& - \int_0^t \pi(s,x) \left( a^{\mathcal{O}}(x, \mathbf{Y}(s)) - \sum_{\tilde{x}} a^{\mathcal{O}}(\tilde{x}, \mathbf{Y}(s)) \pi(s, \tilde{x}) \right) ds \\
& + \sum_{k=1}^{m_1} \int_0^t \left( \frac{\sum_{j \in \mathcal{O}_{\mu_k}} a_j(x - \nu_j', \mathbf{Y}(s^-)) \pi(s^-, x - \nu_j')}{\sum_{\tilde{x}} a^{\mathcal{O}_{\mu_k}}(\tilde{x}, \mathbf{Y}(s^-)) \pi(s^-, \tilde{x})} - \pi(s^-, x) \right) d\tilde{R}_{\mu_k}(s) \qquad \text{(S2)} \\
& \qquad \forall t \geq 0 \text{ and } \forall x \in \mathbb{Z}_{\geq 0}^{n_1} \text{ almost surely.}
\end{aligned}$$

##### S.1.3 Proof of Theorem 1

We first show that the conditional distribution $\pi(t,x)$ satisfies the filtering equation (S2).

**Theorem 1.** *(Validity of the filtering equation). Under the non-explosivity condition* (S1)*, the conditional probability $\pi(t,x)$ (for any $x \in \mathbb{Z}_+^{n_1}$) satisfies* (S2).

*Proof.* Here, we prove the result using the innovation method. First, the non-explosivity condition (S1) suggests that the Chemical Master Equation (CME) precisely characterises the probability of the reaction process $\mathbf{Z}(t)$ [4]. Then, we exploit the CME to find an adequate martingale to represent the conditional distribution as a continuous-time process through the martingale representation theorem. Along these lines, by conditioning the CME w.r.t the observation process $\mathbf{Y}(t)$, we can easily prove that the process

$$\mathcal{M}(t,x) = \pi(t,x) - \left[ \pi(0,x) + \int_0^t \left( \sum_{j=1}^{M} a_j(x - \nu_j', \mathbf{Y}(s)) \pi(s, x - \nu_j') - \sum_{j=1}^{M} a_j(x, \mathbf{Y}(s)) \pi(s,x) \right) ds \right] \qquad \text{(S3)}$$

is a square integrable martingale (due to (S1)) adapted to $\mathcal{Y}_t$. By the martingale representation theorem [5], we can find $\mathcal{Y}_t$-predictable processes $\phi_1(t), \ldots, \phi_{m_1}(t)$ such that this martingale can almost surely be expressed by:

$$\mathcal{M}(t) = \sum_{k=1}^{m_1} \int_0^t \phi_k(s) \left( d\tilde{R}_{\mu_k}(s) - \pi(s^-, a^{\mathcal{O}_{\mu_k}}) ds \right) \qquad \text{(S4)}$$

where $\pi(s^-, a^{\mathcal{O}_{\mu_k}})$ is the left limit of $\pi(s, a^{\mathcal{O}_{\mu_k}}) \triangleq \sum_x a^{\mathcal{O}_{\mu_k}}(x, Y(s))\pi(s,x)$, and $\tilde{R}_{\mu_k}(t) - \int_0^t \pi(s^-, a^{\mathcal{O}_{\mu_k}})ds$ are also $\mathcal{Y}_t$-adaptive martingales. The square integrability of (S3) guarantees the integrability of $\int_0^t (\phi_k(s))^2 \pi(s^-, a^{\mathcal{O}_{\mu_k}})ds$ which ensures that (S4) holds globally. To finalise the proof, we only need to identify $\phi_1(t), \dots, \phi_{m_1}(t)$.

To this end, we can exploit the process $\pi(t,x)e^{-\tilde{R}_{\mu_k}(t)}$ and write its dynamics by

$$\begin{aligned}
&\pi(t,x)e^{-\tilde{R}_{\mu_k}(t)} \\
&= \pi(0,x)e^{-\tilde{R}_{\mu_k}(0)} + \int_0^t e^{-\tilde{R}_{\mu_k}(s)} \left[ \sum_{j=1}^M a_j(x-\nu'_j, \mathbf{Y}(s))\pi(s, x-\nu'_j) - \sum_{j=1}^M a_j(x, \mathbf{Y}(s))\pi(s,x) \right] ds \\
&\quad + \int_0^t e^{-\tilde{R}_{\mu_k}(s^-)} \left[ \sum_{b=1}^{m_1} \phi_b(s) \left( d\tilde{R}_{\mu_b}(s) - \pi(s^-, a^{\mathcal{O}_{\mu_b}})ds \right) \right] \\
&\quad + \int_0^t \pi(s^-, x)(e^{-1}-1)e^{-\tilde{R}_{\mu_k}(s^-)} d\tilde{R}_{\mu_k}(s) + \int_0^t \phi_k(s)(e^{-1}-1)e^{-\tilde{R}_{\mu_k}(s^-)} d[\tilde{R}_{\mu_k}]_s
\end{aligned} \tag{S5}$$

where $[\tilde{R}_{\mu_k}]_t$ is the quadratic variation of $\tilde{R}_{\mu_k}(t)$. Considering the process $\mathbb{1}(\mathbf{X}(t) = x)e^{-\tilde{R}_{\mu_k}(t)}$, we can write:

$$\begin{aligned}
&\mathbb{1}(\mathbf{X}(t) = x)e^{-\tilde{R}_{\mu_k}(t)} \\
&= \mathbb{1}(\mathbf{X}(0) = x)e^{-\tilde{R}_{\mu_k}(0)} + \sum_{j=1}^M \int_0^t \Big[ \mathbb{1}\left(\mathbf{X}(s^-) + \nu'_j = x\right) \left( \mathbb{1}(j \notin \mathcal{O}_{\mu_k}) e^{-\tilde{R}_{\mu_k}(s^-)} + \mathbb{1}(j \in \mathcal{O}_{\mu_k}) e^{-\tilde{R}_{\mu_k}(s^-)-1} \right) \\
&\quad - \mathbb{1}(\mathbf{X}(s^-) = x) e^{-\tilde{R}_{\mu_k}(s^-)} \Big] d\tilde{R}_j(s)
\end{aligned} \tag{S6}$$

where $\tilde{R}_j(t) = R_j\left(\int_0^t a_j(\mathbf{X}(s), \mathbf{Y}(s))ds\right)$.

Then, by denoting $b(s) = \left(e^{-1}-1\right)e^{-\tilde{R}_{\mu_k}(s^-)}$ and applying (S5) and (S6) to the relation $\mathbb{E}\left[\pi(t_2,x)e^{-\tilde{R}_{\mu_k}(t_2)} \middle| \mathcal{Y}_{t_1}\right] = \mathbb{E}\left[\mathbb{1}(\mathbf{X}(t_2) = x)e^{-\tilde{R}_{\mu_k}(t_2)} | \mathcal{Y}_{t_1}\right]$ (for any $t_2 \geq t_1 \geq 0$), we have

$$\begin{aligned}
&\mathbb{E}\left[ -\int_{t_1}^{t_2} b(s) \sum_{j \in \mathcal{O}_{\mu_k}} a_j(x-\nu', \mathbf{Y}(s))ds + \int_{t_1}^{t_2} b(s)\pi(s^-,x)\pi(s^-, a^{\mathcal{O}_{\mu_k}})ds + \int_{t_1}^{t_2} b(s)\phi_k(s)\pi(s^-, a^{\mathcal{O}_{\mu_k}})ds \,\middle|\, \mathcal{Y}_{t_1} \right] \\
&= 0
\end{aligned}$$

for $k = 1, \dots, m_1$ and any $t_2 \geq t_1 \geq 0$. Let us denote a continuous process $\tilde{\mathcal{M}}(t) \triangleq -\int_0^t b(s) \sum_{j \in \mathcal{O}_{\mu_k}} a_j(x-\nu', \mathbf{Y}(s))ds + \int_0^t b(s)\pi(s^-,x)\pi(s^-, a^{\mathcal{O}_{\mu_k}})ds + \int_0^t b(s)\phi_k(s)\pi(s^-, a^{\mathcal{O}_{\mu_k}})ds$, which has finite variation due to the absence of jumps and Brownian motions. The above formula demonstrates that $\tilde{\mathcal{M}}(t)$ is a martingale with respect to $\mathcal{Y}_t$. Therefore, $\tilde{\mathcal{M}}(t)$ is a continuous martingale starting at zero and has a finite total variation; it then follows that this process is almost surely zero [6]. Consequently, for every $k \in \{1, \dots, m_1\}$, every state $x \in \mathbb{Z}_+^{n_1}$, and almost every $t > 0$, it almost surely holds the relation:

$$\pi(t^-, a^{\mathcal{O}_{\mu_k}})\phi_k(t) = \sum_{j \in \mathcal{O}_{\mu_k}} a_j(x-\nu'_j, \mathbf{Y}(t^-))\pi(t^-, x-\nu'_j) - \pi(t^-,x)\pi(t^-, a^{\mathcal{O}_{\mu_k}}).$$

Finally, by inserting the above equality while exploiting the càdlàg property of $\pi(t,x)$ and $\pi(t, a^{\mathcal{O}_{\mu_k}})$ in the martingale representation (S4), we prove the result. □

We then prove the uniqueness of the solution of the filtering equation.

**Theorem 1.** *(Uniqueness).*

*There is a unique (up to indistinguishability) non-negative solution of the filtering equation* (S2) *and such a solution will satisfy*

$$\int_0^t \sum_{x\in\mathbb{Z}_+^{n_1}} \sum_{j=1}^M a_j(x,\mathbf{Y}(s))\pi(s,x)ds < \infty \textit{ almost surely}, \quad \forall t \geq 0. \tag{S7}$$

*Proof.* The existence of the solution is proven in Section S.1.3; here, we only prove the uniqueness of the solution. This proof is not trivial because the propensity functions are not bounded, which invalidates the application of Gronwall's inequality. Instead, we apply an iteration strategy proposed in [4] that successfully proved the uniqueness of certain Chemical Master Equations.

Let $t_1, t_2, \ldots$ be the jumping times of $\mathbf{Y}(t)$ and $t_0 = 0$. Note that if the solution of (S2) is unique between any interval $[t_k, t_{k+1})$ $(k = 0, 1, \ldots)$ given the initial condition at time $t_k$ (i.e., $\pi(t_k, \cdot)$), then the result naturally holds. In the following, we will use the Picard iteration to prove the result.

Between any time interval $[t_k, t_{k+1})$ and any fixed state, we can view (S2) as a linear ordinary differential equation of the form $\dot{y}(t) = -\alpha y(t) + \beta(t)$, with $y(t) = \pi(t, x)$, $\alpha = \sum_{j=1}^M a_j(x, \mathbf{Y}(t_k))$, and

$$\beta(t) = \sum_{j\in\mathcal{U}} a_j(x-\nu_j', \mathbf{Y}(t_k))\pi(t, x-\nu_j') + \pi(t,x)\left(\sum_{x'\in\mathbb{Z}_+^{n_1}} a^{\mathcal{O}}(x', \mathbf{Y}(t_k))\pi(t,x')\right).$$

Therefore, between any time interval $[t_k, t_{k+1})$, the filtering equation (S2) can be equivalently rewritten by

$$\begin{aligned}\pi(t,x) =&\pi(t_k,x)\exp(-\bar{a}(x)t) + \int_{t_k}^t \sum_{j\in\mathcal{U}} \exp(-\bar{a}(x)(t-s))a_j(x-\nu_j', \mathbf{Y}(t_k))\pi(s,x-\nu_j')ds \\ &+ \int_{t_k}^t \exp(-\bar{a}(x)(t-s))\pi(s,x)\left(\sum_{x'\in\mathbb{Z}_+^{n_1}} a^{\mathcal{O}}(x', \mathbf{Y}(t_k))\pi(s,x')\right)ds\end{aligned} \tag{S8}$$

for any $x \in \mathbb{Z}_{\geq 0}^{n_1}$, where $\bar{a}(x) = \sum_{j=1}^M a_j(x, \mathbf{Y}(t_k))$.

Now, we can construct the Picardi iteration through which a solution of (S8) is attained by setting $p^{(0)}(t,z) \equiv 0$ for $t \in [t_k, t_{k+1})$ and

$$\begin{aligned}p^{(\ell+1)}(t,x) =&\pi(t_k,x)\exp(-\bar{a}(x)t) + \int_{t_k}^t \sum_{j\in\mathcal{U}} \exp(-\bar{a}(x)(t-s))a_j(x-\nu_j', \mathbf{Y}(t_k))p^{(\ell)}(s,x-\nu_j')ds \\ &+ \int_{t_k}^t \exp(-\bar{a}(x)(t-s))p^{(\ell)}(s,x)\left(\sum_{x'\in\mathbb{Z}_+^{n_1}} a^{\mathcal{O}}(x', \mathbf{Y}(t_k))p^{(\ell)}(s,x')\right)ds\end{aligned}$$

With mathematical induction, we can check that, for any $t \in [t_k, t_{k+1})$ and $x \in \mathbb{Z}_+^{n_1}$, the succession $\{p^{(\ell)}(t,x)\}_{\ell\in\mathbb{N}}$ is monotonic with respect to $\ell$ and each term of the succession is less than or equal to any non-negative solution of (S8) satisfying (S7). Therefore, as $\ell$ grows to infinity, $p^{(\ell)}(t,x)$ almost surely converges to a non-negative random variable, denoted by $p^{(\infty)}(t,x)$. Moreover, $p^{(\infty)}(t,\cdot)$ is almost surely no greater than any non-negative $\tilde{p}(t,\cdot)$ solving (S8) with initial condition $\pi(t_k,\cdot)$ and satisfying (S7), i.e.,

$$p^{(\infty)}(t,x) \leq \tilde{p}(t,x) \quad \text{for any } t \in [t_k, t_{k+1}) \text{ and any } x \in \mathbb{Z}_+^{n_1}, \text{ almost surely.} \tag{S9}$$

Furthermore, by the convergence of $\left\{p^{(\ell)}(t,x)\right\}_{\ell\in\mathbb{N}}$, we can verify that $p^{(\infty)}(t,\cdot)$ also solves (S8) in the interval $(t_k, t_{k+1})$ with initial condition $\pi(t_k,\cdot)$. Also, from (S9), we can show verify that $p^{(\infty)}(t,x)$ also satisfies (S7) within the integral region $[t_k, t_{k+1})$.

By the dominant convergence theorem, we can prove that any non-negative solution $\tilde{p}(t,\cdot)$ of (S8) satisfying (S7) has a conserved total mass, i.e.,

$$\sum_{x\in\mathbb{Z}_+^{n_1}} p^{(\infty)}(t,x) = \sum_{x\in\mathbb{Z}_+^{n_1}} \pi(t_k,x) = \sum_{x\in\mathbb{Z}_+^{n_1}} \tilde{p}(t,x) \qquad \forall t\in[t_k,t_{k+1}) \text{ almost surely.} \tag{S10}$$

Thanks to (S9) and (S10), all non-negative solutions $\tilde{p}(t,\cdot)$ of (S8) with initial condition $\pi(t_k,\cdot)$ are the same in the time interval $[t_k, t_{k+1})$ almost surely, which proves the result. □

By combining these two theorems, we have proved the existence and the uniqueness of the solution of the filtering equation.

### S.2 Error analysis of the filtered finite state projection (FFSP) algorithms

In this section, we provide more details about the error analysis for the FFSP algorithms. In Section S.2.1, we present the proof of the extended FSP theorem. Then, in Section S.2.2 and Section S.2.3, we show more details about the error analyses for the first and second FFSP algorithms, respectively.

For technical reasons, we only consider the case where the propensity function $a^{\mathcal{O}}(\cdot,y)$ is upper bounded for each fixed $y\in\mathbb{Z}_{\geq 0}^{n_2}$, i.e.,

$$\text{there exists a function } \bar{a}^{\mathcal{O}}(y) \text{ such that } \sup_{x\in\mathbb{Z}_{\geq 0}^{n_1}} a^{\mathcal{O}}(x,y) \leq \bar{a}^{\mathcal{O}}(y). \tag{S11}$$

Moreover, we also assume that the network with only unobserved reactions

$$\tilde{\mathbf{X}}_y(t) = \tilde{\mathbf{X}}_y(0) + \sum_{j\in\mathcal{U}} \nu_j' R_j\left(\int_0^t a_j(\tilde{\mathbf{X}}_y(s),y)ds\right)$$

satisfies the conditions

$$\begin{aligned}&\textstyle\sum_{j=1}^M \mathbb{E}\left[\int_0^t a_j\left(\tilde{\mathbf{X}}_y(s),y\right)ds\right] < \infty \ \forall t\geq 0 \text{ and } \forall y\in\mathbb{Z}_{\geq 0}^{n_2},\\ &\text{as long as } \mathbb{E}\left[\left\|\tilde{\mathbf{X}}_y(0)\right\|_1^q\right] < +\infty \text{ for all } q\geq 0\end{aligned} \tag{S12}$$

which guarantees the process to be non-explosive when initial conditions have finite moments.

#### S.2.1 The extended FSP theorem

**Theorem 3.** *(Extended FSP theorem for un-normalised probability distributions). We consider a set of un-normalised probability distributions $\{\bar{p}(t,\cdot)\}_{t\geq 0}$ defined on a discrete state space $\mathcal{X}=\{x_1,x_2,\ldots\}$ and evolving according to*

$$\dot{\bar{p}}(t,\mathcal{X}) = \mathbb{A}(y)\,\bar{p}(t,\mathcal{X}) \qquad \forall t\geq 0$$

*where $\bar{p}(t,\mathcal{X}) \triangleq \left(\bar{p}(t,x_1),\bar{p}(t,x_2),\ldots\right)^\top$, the variable $y$ is a vector in $\mathbb{Z}_{\geq 0}^{n_2}$, and the matrix $\mathbb{A}(y)$ is defined as the following:*

$$\mathbb{A}_{ij}(y) = \begin{cases} -\sum_{b=1}^M a_b(x_i,y) & \text{if } x_j = x_i \\ \sum_{b\in\mathcal{U}} a_b(x_j,y)\mathbb{1}\left(x_j+\nu_b' = x_i\right) & \text{if } x_j\neq x_i \end{cases}. \tag{S13}$$

*We also consider an FSP system of this infinite dimensional ODE, denoted by* $\{p_{FSP}(t,\cdot)\}_{t\geq 0}$*, which is defined on the same state space but evolves according to*

$$\dot{p}_{FSP}(t,\mathcal{X}) = \left[\begin{array}{c|c} \mathbb{A}_J(y) & \mathbf{0} \\ \hline \mathbf{0} & \mathbf{0} \end{array}\right] p_{FSP}(t,\mathcal{X}) \qquad \forall t \geq 0$$

*where* $p_{FSP}(t,\mathcal{X}) \triangleq (p_{FSP}(t,x_1), p_{FSP}(t,x_2), \dots)^\top$*, the matrix* $\mathbb{A}_J(y)$ *is the first* $J\times J$ *sub-matrix of* $\mathbb{A}(y)$*, and* $p_{FSP}(0,x) = 0$ *for all* $x \notin \mathcal{X}_J$*.*

*Then, under conditions* (S12) *and* $\int_0^t \sum_i |\mathbb{A}_{ii}(y)|\, \bar{p}(s,x_i)ds < +\infty$*, the difference in the* $L_1$ *norm between these two sets of measures can be evaluated by*

$$\|p_{FSP}(t,\mathcal{X}) - \bar{p}(t,\mathcal{X})\|_1 < \epsilon(t) + \|p_{FSP}(0,\mathcal{X}) - \bar{p}(0,\mathcal{X})\|_1 \qquad \forall t \geq 0$$

*where* $\epsilon(t) = \sum_{x\in\mathcal{X}} p_{FSP}(0,x) + \int_0^t \mathbf{1}^\top \mathbb{A}(y)\, p_{FSP}(s,\mathcal{X})ds - \|p_{FSP}(t,\mathcal{X})\|_1$*. (Here,* $\epsilon(t)$ *is computable because the function* $p_{FSP}(t,\cdot)$ *has finite support.)*

*Proof.* The framework of this proof is as follows. First, we introduce an auxiliary process $\hat{p}(t,\mathcal{X})$ which has the same time-evolution dynamics as $\bar{p}(t,\mathcal{X})$ but starts at $p_{\text{FSP}}(0,\mathcal{X})$. Then, the error between $\bar{p}(t,\mathcal{X})$ and $p_{\text{FSP}}(t,\mathcal{X})$ can be bounded by

$$\|\bar{p}(t,\mathcal{X}) - p_{\text{FSP}}(t,\mathcal{X})\|_1 \leq \|\bar{p}(t,\mathcal{X}) - \hat{p}(t,\mathcal{X})\|_1 + \|\hat{p}(t,\mathcal{X}) - p_{\text{FSP}}(t,\mathcal{X})\|_1 \tag{S14}$$

Finally, we prove the result by investigating the two errors on the right hand side of this inequality.

*Construct* $\hat{p}(t,\mathcal{X})$*:* Note that $p_{\text{FSP}}(0,\mathcal{X})$ has a compact support; therefore, the stochastic process $\tilde{\mathbf{X}}_y(t)$ with the initial distribution $\frac{p_{\text{FSP}}(0,\cdot)}{\sum_{\tilde{x}\in\mathcal{X}} p_{\text{FSP}}(0,\tilde{x})}$ is non-explosive due to (S12). By [7, Lemma 1], the process

$$\tilde{p}(t,x) \triangleq \mathbb{E}\left[\mathbb{1}\left(\tilde{\mathbf{X}}_y(t) = x\right) \exp\left(-\int_0^t a^{\mathcal{O}}\left(\tilde{\mathbf{X}}_y(s), y\right) ds\right)\right]$$

evolves according to the same dynamics as $\bar{p}(t,\mathcal{X})$ but starts from $\frac{p_{\text{FSP}}(0,x)}{\sum_{\tilde{x}\in\mathcal{X}} p_{\text{FSP}}(0,\tilde{x})}$. Then, the process $\hat{p}(t,\mathcal{X}) = \tilde{p}(t,\mathcal{X})\left(\sum_{\tilde{x}\in\mathcal{X}} p_{\text{FSP}}(0,\tilde{x})\right)$ satisfies

$$\frac{d}{dt}\hat{p}(t,\mathcal{X}) = \mathbb{A}(y)\hat{p}(t,\mathcal{X}) \quad \forall t \geq 0 \qquad \text{and} \qquad \hat{p}(0,\mathcal{X}) = p_{\text{FSP}}(0,\mathcal{X}).$$

Moreover, by this construction, we can easily check that $\hat{p}(t,\mathcal{X})$ is non-negative component-wise (due to the Metzler matrix $\mathbb{A}(y)$) and has a finite $L_1$ norm at every time point (due to the diagonal dominance of $\mathbb{A}(y)$). Also, by (S12), there holds $\int_0^t \sum_i |\mathbb{A}_{ii}(y)|\, \hat{p}(s,x_i)ds < \infty$.

*Estimate of* $\|\bar{p}(t,\mathcal{X}) - \hat{p}(t,\mathcal{X})\|_1$*:* Let us denote $e_1(t,x_i) = \bar{p}(t,x_i) - \hat{p}(t,x_i)$ for all $x_i \in \mathcal{X}$. Then, by linearity, we have $\dot{e}_1(t,x_i)=\sum_{j=1}^{\infty}\mathbb{A}_{ij}(y)e_1(t,x_j)$ for all $x_i\in\mathcal{X}$ and all $t\geq 0$; furthermore, we also have

$$\begin{aligned}
&\frac{d}{dt^+}\left|e_1(t,x_i)\right| \\
&\triangleq \lim_{dt\to 0^+}\frac{|e_1(t+dt,x_i)|-|e_1(t,x_i)|}{dt} \\
&=\lim_{dt\to 0^+}\frac{\left|e_1(t,x_i)+\sum_{j=1}^{\infty}\mathbb{A}_{ij}(y)e_1(t,x_j)dt+o(dt)\right|-|e_1(t,x_i)|}{dt} \\
&\leq \lim_{dt\to 0^+}\frac{\left|e_1(t,x_i)+\mathbb{A}_{ii}(y)e_1(t,x_i)dt\right|+\left|\sum_{j\neq i}\mathbb{A}_{ij}(y)e_1(t,x_j)dt+o(dt)\right|-|e_1(t,x_i)|}{dt} \\
&\leq \lim_{dt\to 0^+}\frac{\left|e_1(t,x_i)\Big(1+\mathbb{A}_{ii}(y)dt\Big)\right|-|e_1(t,x_i)|}{dt}+\left|\sum_{j\neq i}\mathbb{A}_{ij}(y)e_1(t,x_j)\right| \\
&=\mathbb{A}_{ii}(y)\left|e_1(t,x_i)\right|+\left|\sum_{j\neq i}\mathbb{A}_{ij}(y)e_1(t,x_j)\right| \\
&\leq \sum_{j=1}^{\infty}\mathbb{A}_{ij}(y)\left|e_1(t,x_j)\right| \qquad \forall i\in\mathbb{Z}_{>0} \text{ and } \forall t\geq 0,
\end{aligned}$$

where $\frac{d}{dt^+}$ indicates the right derivative. Thus, we have the expression

$$\left\|\bar{p}(t,\mathcal{X})-\hat{p}(t,\mathcal{X})\right\|_1=\sum_{i=1}^{\infty}|e_1(t,x_i)|\leq\sum_{i=1}^{\infty}|e_1(0,x_i)|+\sum_{i=1}^{\infty}\int_0^t\sum_{j=1}^{\infty}\mathbb{A}_{ij}(y)|e_1(s,x_j)|ds \qquad \forall t\geq 0.$$

Note that the matrix $\mathbb{A}(y)$ is diagonally dominant, so the finiteness of $\int_0^t\sum_i|\mathbb{A}_{ii}(y)|\,\bar{p}(s,x_i)ds$ and $\int_0^t\sum_i|\mathbb{A}_{ii}(y)|\,\hat{p}(s,x_i)ds$ implies the finiteness of the terms $\int_0^t\sum_i\sum_j|\mathbb{A}_{ij}(y)|\,\bar{p}(s,x_j)ds$ and $\int_0^t\sum_i\sum_j|\mathbb{A}_{ij}(y)|\,\hat{p}(s,x_j)ds$. Therefore, by Fubini's theorem, we can further express the error by

$$\begin{aligned}
\left\|\bar{p}(t,\mathcal{X})-\hat{p}(t,\mathcal{X})\right\|_1 &\leq \sum_{i=1}^{\infty}|e_1(0,x_i)|+\int_0^t\sum_{j=1}^{\infty}|e_1(s,x_j)|\underbrace{\left(\sum_{i=1}^{\infty}\mathbb{A}_{ij}(y)\right)}_{\leq 0\text{ component-wise}}ds \\
&\leq \left\|\bar{p}(0,\mathcal{X})-\hat{p}(0,\mathcal{X})\right\|_1
\end{aligned} \tag{S15}$$

where the last line follows from the diagonal dominance of $\mathbb{A}(y)$.

*Estimate of* $\left\|\hat{p}(t,\mathcal{X})-p_{FSP}(t,\mathcal{X})\right\|_1$*:* The analysis in this part is similar to the proof in the classical FSP theorem in [2]. Let us denote $\hat{p}(t,\mathcal{X}_J)\triangleq(\hat{p}(t,x_1),\dots,\hat{p}(t,x_J))^\top$ and $\hat{p}(t,\mathcal{X}_{J'})\triangleq(\hat{p}(t,x_{J+1}),\hat{p}(t,x_{J+2}),\dots)^\top$. Then, we can rewrite the dynamics of $\hat{p}(t,\mathcal{X})$ by

$$\begin{bmatrix}\dot{\hat{p}}(t,\mathcal{X}_J)\\ \dot{\hat{p}}(t,\mathcal{X}_{J'})\end{bmatrix}=\begin{bmatrix}\mathbb{A}_J(y) & \mathbb{A}_{JJ'}(y)\\ \mathbb{A}_{J'J}(y) & \mathbb{A}_{J'J'}(y)\end{bmatrix}\begin{bmatrix}\hat{p}(t,\mathcal{X}_J)\\ \hat{p}(t,\mathcal{X}_{J'})\end{bmatrix} \qquad \forall t\geq 0,$$

where $\mathbb{A}_{JJ}(y)$, $\mathbb{A}_{JJ'}(y)$, $\mathbb{A}_{JJ'}(y)$, and $\mathbb{A}_{J'J'}(y)$ are sub-matrices of $\mathbb{A}(y)$ with proper size; therefore, we can obtain

$$\hat{p}(t,\mathcal{X}_J)=\underbrace{e^{\mathbb{A}_J(y)t}\,p_{\mathrm{FSP}}(0,\mathcal{X}_J)}_{=p_{\mathrm{FSP}}(t,\mathcal{X}_J)}+\int_0^t e^{\mathbb{A}_J(y)\,(t-\tau)}\,\mathbb{A}_{JJ'}(y)\,\hat{p}(\tau,\mathcal{X}_{J'})\,d\tau \qquad \forall t\geq 0$$

where $p_{\mathrm{FSP}}(t, \mathcal{X}_J) = \left(\bar{p}_{\mathrm{FSP}}(t, x_1), \ldots, \bar{p}_{\mathrm{FSP}}(t, x_J)\right)^\top$. Note that $e^{\mathbb{A}_J(y)\,(t-\tau)}$ and $\mathbb{A}_{JJ'}(y)$ are component-wise positive due to the Metzler matrix $\mathbb{A}(y)$; also $\hat{p}(\tau, \mathcal{X}_{J'})$ is component-wise positive due to the discussion in the first part of this proof. Consequently, we can conclude that

$$\hat{p}(t, X_J) \geq p_{\mathrm{FSP}}(t, \mathcal{X}_J) \quad \text{component-wise} \qquad \forall t \geq 0. \tag{S16}$$

Now, we denote the total mass of $\hat{p}(t, X)$ at time $t$ by $c(t) \triangleq \left\|\hat{p}(t, X)\right\|_1$. By the Fubini's theorem, we can express $c(t)$ by

$$\begin{aligned} c(t) &= c(0) + \int_0^t \sum_{j=1}^{\infty} \hat{p}(s, x_j) \underbrace{\left(\sum_{i=1}^{\infty} \mathbb{A}_{i,j}(y)\right)}_{\leq 0 \text{ component-wise}} ds \\ &\leq \sum_{x \in \mathcal{X}} p_{\mathrm{FSP}}(0, x) + \int_0^t \mathbf{1}^\top \mathbb{A}(y)\, p_{\mathrm{FSP}}(s, \mathcal{X}) ds \qquad \forall t \geq 0 \end{aligned}$$

where the last line follows from (S16) and the fact that $p_{\mathrm{FSP}}(t, x) = 0$ for all $x \in \mathcal{X}_{J'}$. Finally, combining this with (S16), we have that

$$\begin{aligned} \left\|\hat{p}(t, \mathcal{X}) - p_{\mathrm{FSP}}(t, \mathcal{X})\right\|_1 &= \sum_{x \in \mathcal{X}} \left[\hat{p}(t, x) - p_{\mathrm{FSP}}(t, x)\right] \\ &= c(t) - \|p_{FSP}(t, \mathcal{X}_J)\|_1 \\ &\leq \sum_{x \in \mathcal{X}} p_{\mathrm{FSP}}(0, x) + \int_0^t \mathbf{1}^\top \mathbb{A}(y)\, p_{\mathrm{FSP}}(s, \mathcal{X}) ds - \|p_{FSP}(t, \mathcal{X})\|_1 \quad \forall t \geq 0. \end{aligned} \tag{S17}$$

By combining (S14), (S15), and (S17), we prove the result. □

##### S.2.2 Error analysis for the first FFSP algorithm

The detailed algorithm for the FFSP method is presented in Algorithm 1.

**Algorithm 1** Filtered finite state projection (FFSP)

**Require:** The observation $\mathbf{Y}(t_k)$ and the initial distribution of the hidden states $\pi(0, x)$.

Initialization: $\pi_{\mathrm{FFSP}}(0, x) = \pi(0, x)$, $\forall x \in \mathcal{X}_J$, and $\pi_{\mathrm{FFSP}}(0, x) = 0$.

**for** $k = 0, 1, 2, \ldots,$ **do**

Approximate the un-normalised filter before the next jump event:

$$\rho_{\mathrm{FFSP}}(t, \mathcal{X}_J) = \exp\left\{\mathbb{A}_J(\mathbf{Y}(t_k))(t - t_k)\right\} \pi_{\mathrm{FFSP}}\left(t_k, \mathcal{X}_J\right) \qquad t \in (t_k, t_{k+1})$$

Normalisation: $\pi_{\mathrm{FFSP}}(t, x) = \frac{\rho_{\mathrm{FFSP}}(t,x)}{\sum_{x \in \mathcal{X}_J} \rho_{\mathrm{FFSP}}(t,x)}$ for all $x \in \mathcal{X}_J$ and $t \in (t_k, t_{k+1})$.

Approximate the filter at the next jump event:

$$\pi_{\mathrm{FFSP}}(t_{k+1}, x) = \frac{\sum_{j \in \mathcal{O}_{k+1}} a_j\left(x - \nu'_j, \mathbf{Y}(t_k)\right) \rho_{\mathrm{FFSP}}\left(t_{k+1}^-, x - \nu'_j\right)}{\sum_{x \in \mathcal{X}_J} \sum_{j \in \mathcal{O}_{k+1}} a_j\left(x, \mathbf{Y}(t_k)\right) \rho_{\mathrm{FFSP}}\left(t_{k+1}^-, x\right)} \qquad x \in \mathcal{X}_J.$$

**end for**

**Theorem 2.** *Let us denote filters* $\pi_{FFSP}(t,\mathcal{X}) \triangleq \left(\pi_{FFSP}(t,x_1),\pi_{FFSP}(t,x_2),\dots\right)^\top$ *and* $\pi(t,\mathcal{X}) \triangleq \left(\pi(t,x_1),\pi(t,x_2),\dots\right)^\top$. *Then, under the conditions* (S1), (S11)*, and* (S12)*, then Algorithm 1 almost surely has an estimation error*

$$\begin{aligned}
&\left\|\pi_{FFSP}(t_k,\mathcal{X})-\pi(t_k,\mathcal{X})\right\|_1 = \epsilon(t_k) && \forall k\in\mathbb{Z}_{\geq 0}\\
&\left\|\pi_{FFSP}(t,\mathcal{X})-\pi(t,\mathcal{X})\right\|_1 = \min\left\{\frac{2\big[\epsilon(t_k)+\tilde{\epsilon}(t)\big]}{\|\rho_{FFSP}(t,\mathcal{X})\|_1},2\right\} && \forall k\in\mathbb{Z}_{\geq 0} \text{ and } \forall t\in(t_k,t_{k+1}),
\end{aligned}$$

*where*

$$\tilde{\epsilon}(t) = \|\rho_{FFSP}(t_k,\mathcal{X})\|_1 + \int_0^t \mathbf{1}^\top \mathbb{A}\left(\mathbf{Y}(t_k)\right)\rho_{FFSP}(s,\mathcal{X})\,ds - \|\rho_{FFSP}(t,\mathcal{X})\|_1$$

*for all* $k\in\mathbb{Z}_{\geq 0}$ *and* $t\in(t_k,t_{k+1})$*, and*

$$\epsilon(t_k) = \begin{cases} \sum_{x\notin\mathcal{X}_J}\pi(t_0,x) & t=0\\ \min\left\{\frac{2\,\bar{a}^{\mathcal{O}_k}\left[\epsilon(t_{k-1})+\tilde{\epsilon}(t_k^-)\right]}{\|\tilde{\rho}_{FFSP}(t_k,\mathcal{X})\|_1},2\right\} & k\in\mathbb{Z}_{>0}\end{cases}$$

*with* $\bar{a}^{\mathcal{O}_k}\triangleq \sup_{x\in\mathbb{Z}^{n_1}_{\geq 0}}\left\{\sum_{j\in\mathcal{O}_k} a_j(x,\mathbf{Y}(t_k))\right\}$.

*Proof.* Here, we prove the result by math induction. Obviously, we have $\|\pi_{\mathrm{FFSP}}(t_0,\mathcal{X})-\pi(t_0,\mathcal{X})\|_1 = \epsilon(t_0)\triangleq\sum_{x\notin\mathcal{X}_J}\pi(t_0,x)$, suggesting that the result holds at time $t_0$. Then, we show that once the result holds at time $t_k$ ($\forall k\in\mathbb{Z}_{>0}$), it also holds in the time interval $(t_k,t_{k+1}]$.

By the extended FSP theorem (whose assumption almost surely holds due to (S1)), we almost surely have

$$\left\|\rho_{\mathrm{FFSP}}(t,\mathcal{X})-\rho(t,\mathcal{X})\right\|_1 \leq \epsilon(t_k)+\tilde{\epsilon}(t) \qquad \forall t\in(t_k,t_{k+1}) \tag{S18}$$

where $\rho_{\mathrm{FFSP}}(t,\mathcal{X}) = \left(\rho_{\mathrm{FFSP}}(t,x_1),\rho_{\mathrm{FFSP}}(t,x_2),\dots\right)^\top$, $\rho(t,\mathcal{X}) = \left(\rho(t,x_1),\rho(t,x_2),\dots\right)^\top$, and

$$\tilde{\epsilon}(t) = \|\rho_{\mathrm{FFSP}}(t_k,\mathcal{X})\|_1 + \int_0^t \mathbf{1}^\top \mathbb{A}\left(\mathbf{Y}(t_k)\right)\rho_{\mathrm{FFSP}}(s,\mathcal{X})\,ds - \|\rho_{\mathrm{FFSP}}(t,\mathcal{X})\|_1.$$

Then, for the normalized filter, we almost surely have that

$$\begin{aligned}
&\left\|\pi_{\mathrm{FFSP}}(t,\mathcal{X})-\pi(t,\mathcal{X})\right\|_1\\
&\leq \left\|\frac{\rho_{\mathrm{FFSP}}(t,\mathcal{X})}{\|\rho_{\mathrm{FFSP}}(t,\mathcal{X})\|_1}-\frac{\rho(t,\mathcal{X})}{\|\rho_{\mathrm{FFSP}}(t,\mathcal{X})\|_1}\right\|_1 + \left\|\frac{\rho(t,\mathcal{X})}{\|\rho_{\mathrm{FFSP}}(t,\mathcal{X})\|_1}-\frac{\rho(t,\mathcal{X})}{\|\rho(t,\mathcal{X})\|_1}\right\|_1\\
&= \frac{\left\|\rho_{\mathrm{FFSP}}(t,\mathcal{X})-\rho(t,\mathcal{X})\right\|_1}{\|\rho_{\mathrm{FFSP}}(t,\mathcal{X})\|_1} + \frac{\Big|\|\rho_{\mathrm{FFSP}}(t,\mathcal{X})\|_1-\|\rho(t,\mathcal{X})\|_1\Big|}{\|\rho_{\mathrm{FFSP}}(t,\mathcal{X})\|_1}\\
&\leq \frac{2\big[\epsilon(t_k)+\tilde{\epsilon}(t)\big]}{\|\rho_{\mathrm{FFSP}}(t,\mathcal{X})\|_1} \qquad \forall t\in(t_k,t_{k+1})
\end{aligned} \tag{S19}$$

where the second line follows from the triangle inequality, and the last line follows from (S18) and the variant triangle inequalities $\|\alpha\|_1-\|\beta\|_1\leq\|\alpha-\beta\|_1$ and $\|\beta\|_1-\|\alpha\|_1\leq\|\alpha-\beta\|_1$.

Now, we analyze the error at the next jump time $t_{k+1}$. We denote

$$\tilde{\rho}(t_{k+1},x) = \sum_{j\in\mathcal{O}_{k+1}} a_j\left(x-\nu_j',\mathbf{Y}(t_k)\right)\rho\left(t_{k+1}^-,x-\nu_j'\right) \qquad \forall x\in\mathbb{Z}^{n_1}_{\geq 0},$$

and

$$\tilde{\rho}_{\text{FFSP}}(t_{k+1}, x) = \sum_{j \in \mathcal{O}_{k+1}} a_j \left(x - \nu_j', \mathbf{Y}(t_k)\right) \rho_{\text{FFSP}} \left(t_{k+1}^-, x - \nu_j'\right) \qquad \forall x \in \mathbb{Z}_{\geq 0}^{n_1}.$$

As before, we denote infinite dimensional vectors $\tilde{\rho}(t_{k+1}, \mathcal{X}) \triangleq \left(\tilde{\rho}(t_{k+1}, x_1), \tilde{\rho}(t_{k+1}, x_2), \dots\right)^\top$, and $\tilde{\rho}_{\text{FFSP}}(t_{k+1}, \mathcal{X}) \triangleq \left(\tilde{\rho}_{\text{FFSP}}(t_{k+1}, x_1), \tilde{\rho}_{\text{FFSP}}(t_{k+1}, x_2), \dots\right)^\top$. Then, by (S18), we almost surely have

$$\|\tilde{\rho}_{\text{FFSP}}(t_{k+1}, \mathcal{X}) - \tilde{\rho}(t_{k+1}, \mathcal{X})\| \leq \bar{a}^{\mathcal{O}_{k+1}} \left[\epsilon(t_k) + \tilde{\epsilon}(t)\right]$$

where $\bar{a}^{\mathcal{O}_{k+1}} \triangleq \sup_{x \in \mathbb{Z}_{\geq 0}^{n_1}} \left\{\sum_{j \in \mathcal{O}_{k+1}} a_j(x, \mathbf{Y}(t_k))\right\}$. Then, similar to the analysis in (S19), the error between the normalized filters is almost surely given by

$$\begin{aligned}
&\|\pi_{\text{FFSP}}(t_{k+1}, \mathcal{X}) - \pi(t_{k+1}, \mathcal{X})\|_1 \\
&\leq \frac{\|\tilde{\rho}_{\text{FFSP}}(t_{k+1}, \mathcal{X}) - \tilde{\rho}(t_{k+1}, \mathcal{X})\|_1}{\|\tilde{\rho}_{\text{FFSP}}(t_{k+1}, \mathcal{X})\|_1} + \frac{\left|\|\tilde{\rho}_{\text{FFSP}}(t_{k+1}, \mathcal{X})\|_1 - \|\tilde{\rho}(t_{k+1}, \mathcal{X})\|_1\right|}{\|\tilde{\rho}_{\text{FFSP}}(t_{k+1}, \mathcal{X})\|_1} \\
&\leq \frac{2\bar{a}^{\mathcal{O}_{k+1}} \left[\epsilon(t_k) + \tilde{\epsilon}(t_{k+1}^-)\right]}{\|\tilde{\rho}_{\text{FFSP}}(t_{k+1}, \mathcal{X})\|_1}
\end{aligned} \tag{S20}$$

By combining (S19), (S20), and the fact that the $L_1$ distance of two probability distribution cannot exceed 2, we prove the result. □

##### S.2.3 Error analysis for the second FFSP algorithm

We present the new algorithm in Algorithm 2 and its estimation error in Section S.2.3.

**Theorem 4.** *Under conditions* (S1) *and* (S11), *Algorithm 2 has the properties that*

- $\pi_{FFSP}(t, x) \leq \pi(t, x)$ *for all* $t \geq 0$ *and* $x \in Z_{\geq 0}^{n_1}$, *and, therefore,*
- *the estimation error is given by* $\|\pi_{FFSP}(t, \mathcal{X}) - \pi(t, \mathcal{X})\|_1 = 1 - \|\pi_{FFSP}(t, \mathcal{X}_J)\|_1$ *for all* $t \geq 0$, *where* $\pi_{FFSP}(t, \mathcal{X}) \triangleq \left(\pi_{FFSP}(t, x_1), \pi_{FFSP}(t, x_2), \dots\right)^\top$ *and* $\pi(t, \mathcal{X}) \triangleq \left(\pi(t, x_1), \pi(t, x_2), \dots\right)^\top$.

*Proof.* The second part of this theorem is a direct consequence of the first one, because with the dominance property, it is very easy to show the result. Therefore, we only prove the first part of this theorem, and we prove it using math induction.

Obviously, the first part of this theorem holds at time $t_0$. Then, we prove that once this result holds at time $t_k$ $(k = 1, 2, \dots)$, it also holds in the time interval $t \in (t_k, t_{k+1}]$. By definition, we almost surely have

$$\begin{aligned}
\rho(t, \mathcal{X}_J) &= e^{\mathbb{A}_J(\mathbf{Y}(t_k))\,(t-t_k)}\, \pi(t_k, \mathcal{X}_J) + \int_{t_k}^{t} e^{\mathbb{A}_J(\mathbf{Y}(t_k))\,(t-s)}\, \mathbb{A}_{JJ'}(\mathbf{Y}(t_k))\, \rho(s, \mathcal{X}_{J'})\, ds \\
&\geq e^{\mathbb{A}_J(\mathbf{Y}(t_k))\,(t-t_k)}\, \pi(t_k, \mathcal{X}_J) \\
&\geq e^{\mathbb{A}_J(\mathbf{Y}(t_k))\,(t-t_k)}\, \pi_{FSP}(t_k, \mathcal{X}_J) \qquad \Big(= \rho_{\text{FFSP}}(t, \mathcal{X}_J)\Big) \qquad \forall t \in (t_k, t_{k+1}).
\end{aligned} \tag{S21}$$

where the inequalities holds element-wisely, and the last two lines follow from the non-negativity of all the terms above. This gives the dominance property between the unnormalized filter and its FFSP counterpart.

**Algorithm 2** Another FFSP algorithm with a tighter error bound only valid under condition (S11)

**Require:** The observation $\mathbf{Y}(t_k)$ and the initial distribution of the hidden states $\pi(0,x)$.

1: Initialization: $\pi_{\text{FFSP}}(0,x) = \pi(0,x)$, $\forall x \in \mathcal{X}_J$.

2: **for** $k = 0, 1, 2, \ldots,$ **do**

3: Approximate the un-normalised filter before the next jump event:

$$\rho_{\text{FFSP}}(t, \mathcal{X}_J) = \exp\left\{\mathbb{A}_J(\mathbf{Y}(t_k))(t - t_k)\right\} \pi_{\text{FFSP}}\left(t_k, \mathcal{X}_J\right) \qquad t \in (t_k, t_{k+1})$$

4: Estimate an upper bound for the normalisation factor before the next jump:

$$c_{\text{FFSP}}(t) = 1 + \int_{t_i}^{t} \mathbf{1}^T \mathbb{A}(\mathbf{Y}(t_k))\, p_{\text{FFSP}}\left(s, \mathcal{X}\right) ds \qquad t \in (t_k, t_{k+1})$$

5: Estimate an upper bound for the normalisation factor at the next jump:

$$\begin{aligned} c_{\text{FFSP}}(t_{k+1}) = &\sum_{x \in \mathcal{X}_J} \sum_{j \in \mathcal{O}_{k+1}} a_j\left(x, \mathbf{Y}(t_k)\right) \rho_{\text{FFSP}}(t_{k+1}^-, x) \\ &+ \bar{a}^{\mathcal{O}_{k+1}} \left[c_{\text{FFSP}}(t_{k+1}^-) - \left\|\rho_{\text{FFSP}}(t_{k+1}^-, \mathcal{X})\right\|_1\right] \end{aligned}$$

with $\bar{a}^{\mathcal{O}_{k+1}} \triangleq \sup_{x \in \mathbb{Z}_{\geq 0}^{n_1}} \left\{\sum_{j \in \mathcal{O}_{k+1}} a_j(x, \mathbf{Y}(t_k))\right\}$ whose existence is provided by (S11).

6: Normalisation: for every $x \in X_J$, we compute

$$\begin{aligned} \pi_{\text{FFSP}}(t,x) = &\frac{\rho_{\text{FFSP}}(t,x)}{c_{\text{FFSP}}(t)} \qquad t \in (t_k, t_{k+1}) \\ \pi_{\text{FFSP}}(t_{k+1},x) = &\frac{\sum_{j \in \mathcal{O}_{k+1}} a_j\left(x - \nu_j', y(t_k)\right) \rho_{\text{FFSP}}\left(t_{k+1}^-, x - \nu_j'\right)}{c_{\text{FFSP}}(t_{k+1})}. \end{aligned}$$

7: **end for**

Then, we estimate the normalization factors. For every $t \in (t_k, t_{k+1})$, there almost surely holds

$$\begin{aligned}
\|\rho(t,\mathcal{X})\|_1 &= \|\pi(t_k,\mathcal{X})\|_1 + \int_{t_k}^{t} \mathbf{1}^\top \mathbb{A}(\mathbf{Y}(t_k))\rho(s,\mathcal{X})ds \qquad \text{(by definition \& Fubini's theorem)}\\
&\leq 1 + \int_{t_k}^{t} \mathbf{1}^\top \mathbb{A}(\mathbf{Y}(t_k))\rho_{\mathrm{FFSP}}(s,\mathcal{X})ds\\
&= c_{\mathrm{FFSP}}(t)
\end{aligned} \tag{S22}$$

where the second line follows from (S21) and the non-positivity of $1^\top \mathbb{A}(\mathbf{Y}(t_k))$ (c.f. its definition). Also, we have that

$$\begin{aligned}
&\sum_{x\in\mathbb{Z}_{\geq 0}^{n_1}} \sum_{j\in\mathcal{O}_{k+1}} a_j(x,\mathbf{Y}(t_k))\rho(t_{k+1}^-,x)\\
&= \sum_{x\in\mathbb{Z}_{\geq 0}^{n_1}} \sum_{j\in\mathcal{O}_{k+1}} a_j(x,\mathbf{Y}(t_k))\rho_{\mathrm{FFSP}}(t_{k+1}^-,x)\\
&\quad + \sum_{x\in\mathbb{Z}_{\geq 0}^{n_1}} \left[\sum_{j\in\mathcal{O}_{k+1}} a_j(x,\mathbf{Y}(t_k))\right]\left[\rho(t_{k+1}^-,x) - \rho_{\mathrm{FFSP}}(t_{k+1}^-,x)\right]\\
&\leq \sum_{x\in\mathcal{X}_J} \sum_{j\in\mathcal{O}_{k+1}} a_j(x,\mathbf{Y}(t_k))\rho_{\mathrm{FFSP}}(t_{k+1}^-,x) + a^{\mathcal{O}_{i+1}}\left[c_{\mathrm{FFSP}}(t_{k+1}^-) - \left\|\rho_{\mathrm{FFSP}}(t_{k+1}^-,\mathcal{X})\right\|_1\right]\\
&= c_{\mathrm{FFSP}}(t_{k+1})
\end{aligned} \tag{S23}$$

where the inequality follows from (S21) and (S11). Finally, the normalized filter almost surely satisfies

$$\pi(t,x) = \frac{\rho(t,x)}{\|\rho(t,\mathcal{X})\|_1} \geq \frac{\rho_{\mathrm{FFSP}}(t,x)}{c_{\mathrm{FFSP}}(t)} = \pi_{\mathrm{FFSP}}(t,x) \qquad \forall t \in (t_k,t_{k+1}), \qquad \text{(due to (S21) and (S22))}$$

and

$$\pi(t_{k+1},x) = \frac{\rho(t_{k+1},x)}{\sum_{x\in\mathbb{Z}_{\geq 0}^{n_1}} \sum_{j\in\mathcal{O}_{k+1}} a_j(x,\mathbf{Y}(t_k))\rho(t_{k+1}^-,x)} \geq \frac{\rho_{\mathrm{FFSP}}(t_{k+1},x)}{c_{\mathrm{FFSP}}(t_{k+1})} = \pi_{\mathrm{FFSP}}(t_{k+1},x), \qquad \text{(due to (S21) and (S23))}$$

which suggests that the first part of the theorem holds in $(t_k, t_{k+1}]$. Therefore, we prove this theorem. □

### S.3 Derivation of the Kalman and Extended Kalman Filters for the Chemical Langevin Equation

In this section, we focus on the derivation of the Kalman and Extended Kalman filters using the Chemical Langevin Equation as the hidden model dynamics.

We examine an intracellular chemical reaction system consisting of $n$ species $(S_1, \ldots, S_n)$ and $M$ reactions:

$$\sum_{i=1}^{n} \xi_{ik} S_i \xrightarrow{c_k} \sum_{i=1}^{n} \xi'_{ik} S_i, \quad k = 1, \ldots, M,$$

where $\xi_{ik}$ and $\xi'_{ik}$ represent the quantities of $S_i$ molecules consumed and produced in the $k^{\text{th}}$ reaction, respectively. We denote the propensity functions, representing the rates of these $M$ reactions, as

$a_1, a_2, \ldots, a_M$. Furthermore, we define the stoichiometry vector associated with the $k^{\text{th}}$ reaction channel as $\nu_k \triangleq \xi'_k - \xi_k$ and the stoichiometry matrix as:

$$\mathbb{S} = \begin{bmatrix} | & & | \\ s_1 & \ldots & s_M \\ | & & | \end{bmatrix}.$$

In the low copy number regime, the interactions among intracellular biomolecular species are inherently stochastic, and their dynamics are typically modeled by a stochastic dynamic equation known as the Random Time Change (RTC) representation [4]:

$$\mathbf{Z}(t) = \mathbf{Z}(0) + \sum_{j=1}^{M} s_j R_j \left( \int_0^t a_j(\mathbf{Z}(s))\, ds \right). \tag{S24}$$

Here, $\mathbf{Z}(t) \in \mathcal{Z} \subseteq \mathbb{Z}^n_{\geq 0}$ is a continuous-time discrete-state Markov chain (CTMC) that tracks the copy numbers of molecules, and $R_1, \ldots, R_M$ are counting processes that keep track of the occurrence of each reaction event $j = 1, \ldots, M$ until time $t > 0$. If $\Omega$ is the volume of the system, then we can define another CTMC $\left\{ \mathbf{Z}^\Omega(t) | t \geq 0 \right\}$ by $\mathbf{Z}^\Omega(t) = \frac{\mathbf{Z}(t)}{\Omega}$, which gives the concentrations of all the $n$ species at time $t$. If for each $j = 1, \ldots, M$, we build a function $\tilde{a}_j$ such that $a_j(\Omega \mathbf{Z}^\Omega(t)) \approx \Omega \tilde{a}_j(\mathbf{Z}^\Omega(t))$, then we can write:

$$\mathbf{Z}^\Omega(t) = \mathbf{Z}^\Omega(0) + \frac{1}{\Omega} \sum_{j=1}^{M} s_j R_j \left( \int_0^t \Omega \tilde{a}_j(\mathbf{Z}^\Omega(s))\, ds \right). \tag{S25}$$

Thus, $\mathbf{Z}^\Omega(t) \approx \frac{\mathbf{Z}(t)}{\Omega}$. We define $\tilde{\mathbf{Z}}(t)$ to be the solution of the following stochastic differential equation:

$$d\tilde{\mathbf{Z}}(t) = \mathbb{S}\tilde{\mathbf{A}}(\tilde{\mathbf{Z}}(t))dt + \mathbb{S}\mathbb{D}(\tilde{\mathbf{Z}}(t))d\mathbf{W}(t), \tag{S26}$$

where $\tilde{\mathbf{A}}(\tilde{\mathbf{Z}}(t)) = \begin{bmatrix} \tilde{a}_1(\tilde{\mathbf{Z}}(t)) \\ \tilde{a}_2(\tilde{\mathbf{Z}}(t)) \\ \vdots \\ \tilde{a}_M(\tilde{\mathbf{Z}}(t)) \end{bmatrix}$, $\mathbb{D}(\tilde{\mathbf{Z}}(t)) = \left( \mathbb{D}_{ii}(\tilde{\mathbf{Z}}(t)) \right)_{i=1,\ldots,M} = \sqrt{\frac{\tilde{a}_i(\tilde{\mathbf{Z}}(t))}{\Omega}}$, and $\mathbf{W}(t) \in \mathbb{R}^M$ is a vector-valued standard Brownian motion. Then $\tilde{\mathbf{Z}}(t)$ is defined to be the **diffusion approximation** or the **Langevin approximation** [8] for $\mathbf{Z}^\Omega(t)$. In particular, $\tilde{\mathbf{Z}}(t) \approx \mathbf{Z}^\Omega(t)$ for large values of $\Omega$.

We now aim to construct the Kalman and Extended Kalman filters using the diffusion approximation as a model for the hidden dynamics. We assume that $\tilde{\mathbf{Z}}(t)$ represents the hidden system, with part of this hidden system being observed through the process $Y^\Omega(t) \in \mathbb{R}$. Specifically, the full hidden and observation model can be written as:

$$\begin{cases} d\tilde{\mathbf{Z}}(t) = \mathbb{S}\tilde{\mathbf{A}}(\tilde{\mathbf{Z}}(t))dt + \mathbb{S}\mathbb{D}(\tilde{\mathbf{Z}}(t))d\mathbf{W}(t), \\ dY^\Omega(t) = h(\tilde{\mathbf{Z}}(t))dt + \rho dV(t), \end{cases} \tag{S27}$$

where $\rho \in \mathbb{R}$ is the observation variance noise and $V(t) \in \mathbb{R}$ is a standard Brownian motion. Assuming we perturb the dynamical system around the deterministic process $\bar{\mathbf{Z}}(t)$:

$$\begin{cases} \frac{d\bar{\mathbf{Z}}(t)}{dt} = \mathbb{S}\tilde{\mathbf{A}}(\bar{\mathbf{Z}}(t)), \\ \bar{\mathbf{Z}}(0) = \mathbf{Z}_0, \end{cases} \tag{S28}$$

and that the perturbations from this system are small, we can apply the Kalman filter to the linearized dynamical system [2]. The Kalman filter evolution equations for the estimated conditional mean $\hat{\mathbf{Z}}(t)$ and conditional covariance matrix $\mathbb{R}(t)$ are as follows:

$$\begin{cases} d\hat{\mathbf{Z}}(t) = \left[\mathbb{SJ}\left(\bar{\mathbf{Z}}(t)\right)\hat{\mathbf{Z}}(t) + \mathbb{S}\tilde{\mathbf{A}}\left(\bar{\mathbf{Z}}(t)\right) - \mathbb{SJ}\left(\bar{\mathbf{Z}}(t)\right)\bar{\mathbf{Z}}(t)\right] dt \\ \qquad + \frac{\mathbb{R}(t)h'^T(\bar{\mathbf{Z}}(t))}{\rho^2}\left[dY^\Omega - \left(h'(\bar{\mathbf{Z}}(t))\hat{\mathbf{Z}}(t) + h(\bar{\mathbf{Z}}(t)) - h'(\bar{\mathbf{Z}}(t))\bar{\mathbf{Z}}(t)\right)\right], \\ \frac{d\mathbb{R}(t)}{dt} = \mathbb{SJ}\left(\bar{\mathbf{Z}}(t)\right)\mathbb{R}(t) + \mathbb{R}(t)\mathbb{J}^T\left(\bar{\mathbf{Z}}(t)\right)\mathbb{S}^T + \mathbb{SDD}^T(\bar{\mathbf{Z}}(t))\mathbb{S}^T - \frac{\mathbb{R}(t)h'^T(\bar{\mathbf{Z}}(t))h'(\bar{\mathbf{Z}}(t))}{\rho^2}, \end{cases} \tag{S29}$$

where $\mathbb{J}\left(\bar{\mathbf{Z}}(t)\right)$ is the Jacobian of the propensities evaluated at $\bar{\mathbf{Z}}(t)$.

If we linearize the hidden and observed dynamics around the conditional estimated mean $\hat{\mathbf{Z}}(t)$ instead, we can then apply the Extended Kalman filter, whose evolution equations for the conditional mean and variance are:

$$\begin{cases} d\hat{\mathbf{Z}}(t) = \mathbb{SJ}\left(\hat{\mathbf{Z}}(t)\right) dt + \frac{\mathbb{R}(t)h'^T(\hat{\mathbf{Z}}(t))}{\rho^2}\left[dY^\Omega - h(\hat{\mathbf{Z}}(t))\right], \\ \frac{d\mathbb{R}(t)}{dt} = \mathbb{SJ}\left(\hat{\mathbf{Z}}(t)\right)\mathbb{R}(t) + \mathbb{R}(t)\mathbb{J}^T\left(\hat{\mathbf{Z}}(t)\right)\mathbb{S}^T + \mathbb{SDD}^T(\hat{\mathbf{Z}}(t))\mathbb{S}^T - \frac{\mathbb{R}(t)h'^T(\hat{\mathbf{Z}}(t))h'(\hat{\mathbf{Z}}(t))}{\rho^2}. \end{cases} \tag{S30}$$

##### S.3.1 Nonlinear Network Example

We have tested the Kalman and Extended Kalman filters on the nonlinear network with feedback, as shown in Fig. S1, and used the Bootstrap Particle Filter in [7] and the Filtered State Projection Method (FFSP) as benchmarks of validation. As standard practice in the framework of the FFSP, we have a continuous time discrete state Markov Chain (CTMC) $\mathbf{Z}(t) = (Z_1, Z_2, X_1) = (z_1, z_2, x_1) \in \mathbb{Z}^3_{\geq 0}$ associated with the chemical reacting system, as depicted in panel a) of Fig. S1. Moreover, we further decompose the network in hidden species and observed species denoted with $\mathbf{X}(t)$ and $\mathbf{Y}(t)$ respectively.

As shown in panel (a) of Fig. S1, we assume that $\mathbf{X}(t) = (Z_1(t), Z_2(t)) = (z_1, z_2) \in \mathbb{Z}^2_{\geq 0}$ constitutes the hidden species of the network, which are being produced and jointly degraded. Together, they catalytically influence the production of $\mathbf{Y}(t) = (X_1(t)) = (x_1) \in \mathbb{Z}_{\geq 0}$, the observed species in the network.

In panel (b) of Fig. S1, an SSA trajectory of $X_1$ has been generated to serve as the observation process trajectory for the different filters. Panel (c) of Fig. S1 shows the network's reactions and the nonlinear feedback exerted by $Z_1$ and $Z_2$ on $X_1$. Panel (d) shows a comparison of the mean and variance of each of the species of the network computed with the SSA and the diffusion approximation in Eq. (S26).

For the species $Z_1$ and $Z_2$, the two methods show a strong agreement in estimating the unconditional statistics; however, for the species $X_1$, a slight mismatch can be seen in the estimation of the mean, and a much higher variance is estimated by the diffusion approximation approach. Such high values for the variance might be due to the fact that, for this choice of kinetic parameters, the feedback exerted by $Z_1$ and $Z_2$ on $X_1$ is highly nonlinear and may produce a non-Gaussian type of noise, which contrasts with the assumptions of the diffusion approximation framework.

The following propensities functions have been adopted: $a_1(z_1, z_2, x_1) = c_1, a_2(z_1, z_2, x_1) = c_2, a_3(z_1, z_2, x_1) = c_3 z_1 z_2, a_4(z_1, z_2, x_1) = c_4 + \frac{k3\left(\frac{z_1}{k_1}\right)}{k_4 + \left(\frac{z_1}{k_1}\right) + \left(\frac{z_2}{k_2}\right)}, a_5(z_1, z_2, x_1) = c_5 x_1$. In order to apply the Kalman and Extended Kalman filters, we employed the following rescaling of the reaction rates parameters: $\tilde{c}_1 = \frac{c_1}{\Omega}, \tilde{c}_2 = \frac{c_2}{\Omega}, \tilde{c}_3 = c_3\Omega, \tilde{c}_4 = \frac{c_4}{\Omega}, \tilde{c}_5 = c_5, \tilde{k}_1 = \frac{k_1}{\Omega}, \tilde{k}_2 = \frac{k_2}{\Omega}, \tilde{k}_3 = \frac{k_3}{\Omega}, \tilde{k}_4 = k_4$. This rescaling allows to write

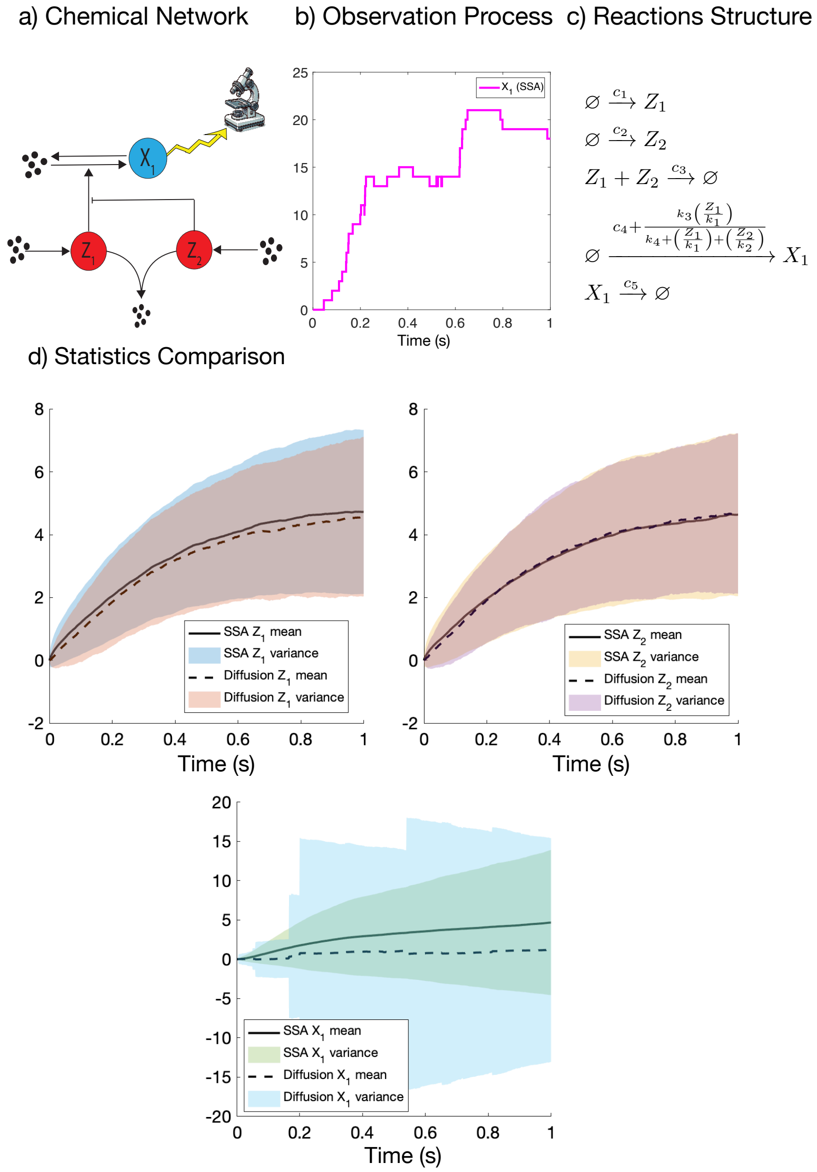


**Fig. S1: Non-Linear Network with Feedback.**
**a) Chemical Reaction Network Structure:** The hidden species $Z_1$ and $Z_2$, highlighted in red, are produced and jointly degraded. They also catalytically influence the production of $X_1$, the observed species in the network.
**b) Observation SSA Process Trajectory:** The trajectory of species $X_1$ is used as input to the filters.
**c) Chemical Reactions Structure:** The network's chemical reactions are structured with the following parameters: $c_1 = 10$, $c_2 = 10$, $c_3 = 0.5$, $c_4 = 0$, $c_5 = 0.5$, $k_1 = 1000$, $k_2 = 1$, $k_3 = 1000$, $k_4 = 0.04$. The initial condition is $[z_{10}, z_{20}, x_{10}]^T = [0, 0, 0]^T$.
**d) Comparison of SSA and Chemical Langevin Equation Statistics.**
This figure compares the statistics of the non-linear network using the Stochastic Simulation Algorithm (SSA) and the Chemical Langevin Equation (CLE). The comparison is based on statistics computed from 1000 trajectories generated by the SSA and simulations of the diffusion approximation defined in Eq. (S26). With this choice of reaction rate parameters, we observe a strong correlation in the statistics of $Z_1$ and $Z_2$, and a slight mismatch in the mean values of $X_1$. The diffusion approximation yields a higher variance, likely due to the highly non-linear feedback exerted by $Z_1$ and $Z_2$. This feedback produces noise that deviates from unimodal Gaussian distributions, which is assumed in the diffusion approximation.

$a_j(\Omega \mathbf{Z}^{\Omega}(t)) \approx \Omega \tilde{a}_j(\mathbf{Z}^{\Omega}(t))$, where $\mathbf{Z}^{\Omega}(t)$ is the CTMC keeping track of the concentration of all the species at time $t > 0$ satisfying Eq. (S25). In particular, in the hidden and observed model in Eq. (S27), $\tilde{\mathbf{Z}}(t) = \left(\tilde{Z}_1(t), \tilde{Z}_2(t), \tilde{X}_1(t)\right)$ and $h\left(\tilde{\mathbf{Z}}(t)\right) = \tilde{X}_1(t)$.

In Fig. S2, the filter estimates of the conditional expectations and standard deviations of the hidden processes $Z_1$ and $Z_2$ are reported for the Kalman and Extended Kalman filters, Bootstrap Particle Filters (BPF), and the Filtered Finite State Projection (FFSP) method for different values of the observation noise parameter $\rho$ from Eq. (S27). While BPF and FFSP assume a noise-free observation model, directly taking the SSA trajectory as input to the filtering algorithm, the Kalman and Extended Kalman filters are built on a noisy observation model, as detailed in Eq. (S27), to derive the filtering equations.

To compare the filters on a noise-free type of observation model, we fed the Kalman and Extended Kalman filters in Eq. (S29) and Eq. (S30) with the exact SSA trajectory of $X_1$ shown in panel b) of Fig. S1, choosing low values of the observation noise parameter $\rho$. The filter results shown in Fig. S2 for the different values of $\rho$ reveal appreciable similarities, highlighting consistency with the noise-free observation model assumption. While the BPF and FFSP exhibit good performance in reproducing the hidden process dynamics, the Kalman and Extended Kalman filters show accentuated discrepancies, likely due to the non-linearities of the model and the lack of Gaussianity for this choice of kinetic parameters.

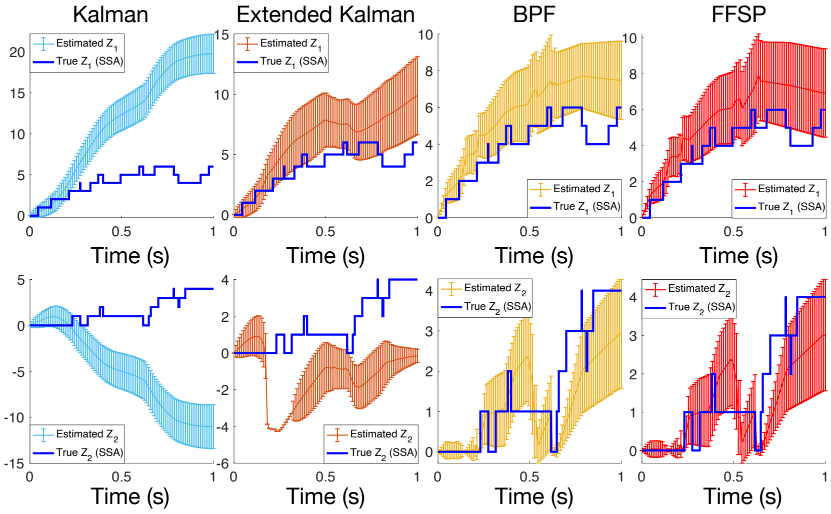


$a)\,\rho = 0.02$

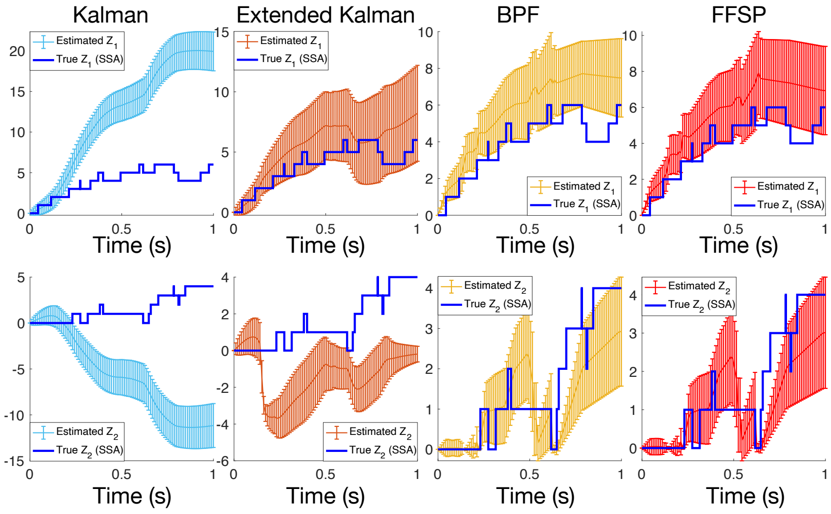


$b)\,\rho = 0.015$

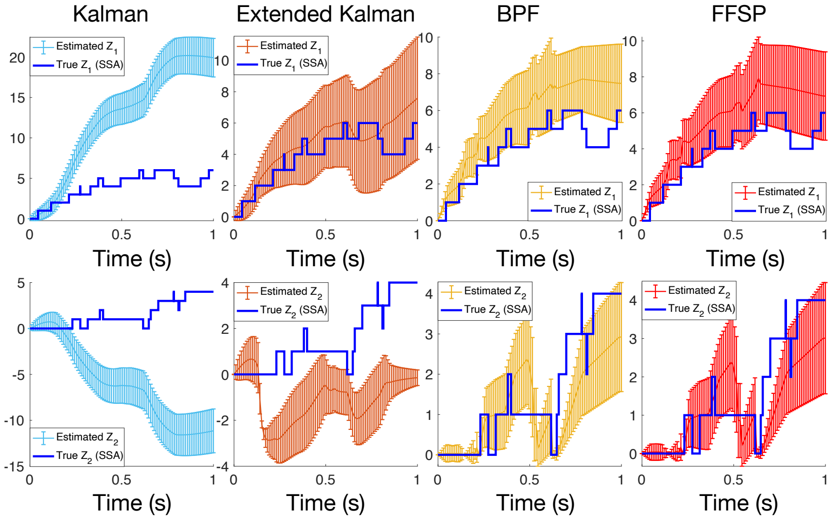


$c)\,\rho = 0.013$

**Fig. S2: Non-linear Network Example: Filter estimates for different values of the observation noise parameter $\rho$ from the observation model defined in Eq. (S27).**
Panels (a)-(c) show the hidden SSA trajectories of processes $Z_1$ and $Z_2$ that generated the observation process trajectory in Fig. S1. The observation process trajectory from panel (b) in Fig. S1 was fed to the Kalman and Extended Kalman filters, Bootstrap Particle Filter (BPF), and Filtered Finite State Projection Method (FFSP). The conditional expectations and standard deviation estimates by the filters are shown in all three panels against the exact SSA trajectories of the hidden processes $Z_1$ and $Z_2$.
